# TaxaAudit: A statistical model of consensus in phylogenetics identifies non-monophyletic taxa

**DOI:** 10.64898/2026.11.09.765192

**Authors:** John B. Allard, Jack M. Craig, Sudhir Kumar

## Abstract

Interpreting variation in genes and traits across species requires understanding their evolutionary relationships. Taxonomic hierarchies provide a practical approximation of the tree of life, with the NCBI taxonomy linking this framework directly to public sequence data. Some named groups, however, are not evolutionary clades, and recognizing these exceptions requires knowledge scattered across the phylogenetics literature. Here we introduce TaxaAudit, a statistical model that identifies taxonomic groups whose monophyly is repeatedly contradicted by published phylogenies. The model accounts for background taxonomic conflict and the ability of each study’s species sampling to expose non-monophyly. Applied to 5,865 molecular phylogenies from TimeTree, TaxaAudit flagged 516 NCBI taxa at family rank and below for significant recurrent contradiction of monophyly. Excluding these taxa’s monophyly requirements improved the taxonomy’s agreement with independent phylogenomic reference trees and phylogenies withheld from discovery. Flagged taxa were also 3.61 times as likely as matched controls to have their monophyly challenged in publications outside the discovery set. In tree-inference experiments, excluding these monophyly constraints improved agreement with reference phylogenies relative to enforcing the unmodified taxonomy. By identifying groups whose monophyly should not be assumed, TaxaAudit lets researchers draw on published phylogenetic knowledge without reviewing the specialist literature for every taxon. This supports broader use of the NCBI taxonomy as an evolutionary framework for biological comparisons.

## Introduction

Evolutionary relationships provide the context for interpreting biological similarities and differences across species. Although explicit phylogenies are available for many groups of organisms, taxonomic hierarchies offer a readily accessible approximation of the tree of life. Their nested groups provide a starting point for identifying related species and choosing comparisons at different evolutionary depths.

The NCBI Taxonomy is a hierarchical classification of organisms used by GenBank and the other databases of the International Nucleotide Sequence Database Collaboration to organize public nucleotide and protein sequence records (Federhen 2012; Schoch et al. 2020). Its stable numerical identifiers maintain connections among taxonomic names and molecular records as classifications change. Researchers use this hierarchy to retrieve sequences and define study groups (Bennett et al. 2018), organize orthology searches (Kriventseva et al. 2015), and partition large phylogenetic analyses (Smith et al. 2009). It also supports metagenomic sequence classification (Wood and Salzberg 2014) and guides phylogenomic reconstruction (Fang et al. 2013). NCBI groups define evolutionary levels for hierarchical orthology inference (Kuznetsov et al. 2023; Hernández-Plaza et al. 2023) and are used to reconstruct gene-family origins (Barrera-Redondo et al. 2023). Phylogeny synthesis resources, including TimeTree of Life and Open Tree of Life, have likewise used the NCBI taxonomy or taxonomies derived partly from it (Hinchliff et al. 2015; Kumar et al. 2022). Compared with broader species checklists such as Catalogue of Life (Hobern et al. 2021), this integration with molecular databases makes the NCBI Taxonomy particularly useful for computational research. The NCBI Taxonomy publication received almost 1,000 citations in 2026 alone ([Google Scholar](https://scholar.google.com/)), illustrating how widely this infrastructure is used in biological research.

Yet the utility of taxonomy as an evolutionary framework depends on whether named taxa correspond to clades. Taxonomy changes for several reasons. New phylogenetic and morphological evidence can lead to the resurrection of previously synonymized genera and the reassignment of species, as in recent revisions of Corydoradinae (Dias et al. 2025). These taxonomic actions are intended to correct discrepancies between the taxonomy and evolutionary history, but they can expose further cases where a recognized taxon does not represent a single evolutionary lineage. NCBI explicitly cautions that its taxonomy is not an authoritative source for nomenclature or classification (National Center for Biotechnology Information, n.d.). Taxonomic disagreements persist even among well-studied groups, as illustrated by efforts to reconcile global bird checklists through AviList (Rheindt et al. 2025). Researchers and computational workflows that treat every NCBI group as monophyletic may therefore unknowingly inherit evolutionary assumptions that conflict with credible phylogenetic evidence.

Baleen whales provide a clear example. The NCBI and other taxonomies recognize the humpback whale as a member of the monotypic genus *Megaptera* and the gray whale, *Eschrichtius robustus*, in a separate monotypic family. However, molecular phylogenetic analyses nearly universally place both within the clade containing all recognized species of *Balaenoptera* (Árnason et al. 2018; McGowen et al. 2020). Therefore, the continued recognition of *Megaptera* and *Eschrichtius* by most working taxonomists renders *Balaenoptera* polyphyletic. A researcher familiar with cetacean taxonomy would recognize this exception; a general-purpose computational pipeline would not.

Comparable cases are scattered across the taxonomic and systematic literature, but recognizing them depends on taxon-specific knowledge that is difficult to apply consistently across the tree of life. Overcoming this challenge by simply counting conflicting trees is inadequate because publications differ in species sampling, phylogenetic resolution, and background taxonomic conflict. A tree lacking the humpback whale, for example, may recover *Balaenoptera* as monophyletic without revealing the problem. Furthermore, in less clear-cut cases, the sparse published record and high background levels of taxonomic discord make it difficult to reproducibly identify real taxonomic problems.

Here we introduce TaxaAudit, a statistical model that identifies taxonomic groups whose monophyly is repeatedly contradicted in a sample of published phylogenies more often than expected from background conflict, while accounting for species sampling and excluding unresolved observations. Comparing against the richest sample of timed molecular phylogenies available, 5,865 trees from the TimeTree of Life database, allowed us to flag 516 NCBI taxa at family rank or below for significant recurrent contradiction of monophyly after multiple-testing correction. We evaluated these findings using withheld publications, phylogenomic reference trees, and molecular tree-inference experiments. The resulting catalog of flagged taxa identifies taxa whose monophyly should not be assumed in downstream analyses and provides candidates for examination and potential revision by taxonomists (Fig. 1A).

**Figure 1.**
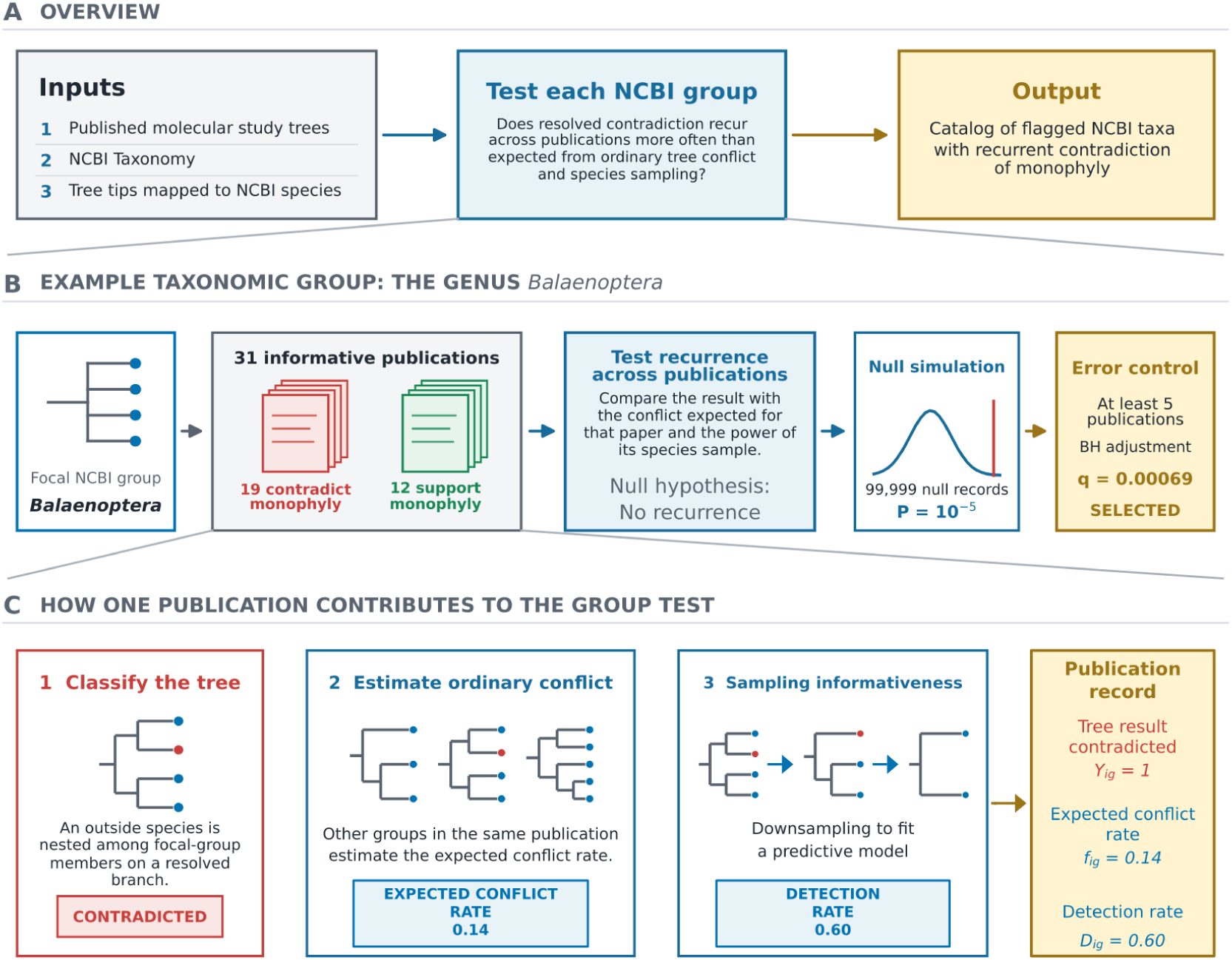
Statistical identification of NCBI groups that are repeatedly contradicted by published phylogenies. **(A)** Overview of the complete analysis, from molecular study trees and the NCBI Taxonomy to a catalog of flagged NCBI taxa whose monophyly is repeatedly contradicted. **(B)** The genus Balaenoptera is followed through the procedure using its observed data: 19 of 31 informative publications contradicted its monophyly, producing a parametric-bootstrap *P* = 10^−5^ and an FDR-adjusted *q* = 0.00069. **(C)** Construction of one publication-level record. Its support or contradiction of monophyly is considered together with the ordinary conflict rate estimated from other groups of the same rank in that publication and the probability that its species sampling could expose a violation. Sparse publication-level estimates of ordinary conflict are moderated toward the overall rate, and sampling informativeness is estimated using a monotone gradient boosted tree model trained on examples generated by subsampling trees with observed conflicts. A tree that is compatible with monophyly but unresolved contributes no outcome.

## Results

### Statistical identification of taxonomic groups contradicted by published phylogenies

We interrogated the NCBI Taxonomy (downloaded 22 July 2026) in comparison with 5,865 molecular tree files from the TimeTree of Life database (Kumar et al. 2022). TimeTree provides an expert-curated set of timed molecular phylogenies with explicit published provenance, making it an ideal test set, but a researcher could alternatively provide their own set of trees, timed or untimed, using molecular data or any other type, to facilitate their own hypothesis testing. We evaluated taxa at family rank and below, including named unranked clades, for contradictions of monophyly in published trees. Administrative or placeholder nodes were not tested as biological groups, but their descendants remained eligible. The study trees represented 26,291 NCBI taxa in this range. Of these, 18,109 had at least one informative publication, yielding 73,181 informative assessments of individual taxa within publications.

For every group represented by at least two species in a study tree, we asked whether its members were recovered as monophyletic, positively contradicted by a resolved split, or compatible with monophyly but unresolved (e.g., participated in a polytomy with outside species, Fig. 1C). The latter state supplied no evidence for or against the group. We then compared the observed record with a background probability of random apparent non-monophyly learned from other groups of the same rank in the same publication and adjusted for the number of focal-group species sampled. For publication *i* and group *g*, *Y_ig_*was 1 when the publication contradicted the group’s monophyly and 0 when it recovered the group as monophyletic. We modeled the probability of contradiction as

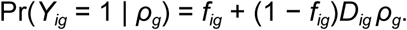

Here, *f_ig_* is the expected probability of background conflict for group *g* in publication *i*, *D_ig_* is the probability that the publication’s species sampling would expose a real violation, and *ρ_g_*is the recurrence strength fitted for group *g* and shared across all of its informative publications. Under the null hypothesis *ρ_g_* = 0, contradiction occurs only at the estimated background rate. We obtained a likelihood-ratio *P* value by simulating publication outcomes under this null model, refitting *ρ_g_* to each simulated data set, and measuring how often the simulated likelihood-ratio statistic was at least as large as the observed statistic (Fig. 1B,C; see Methods for details).

Many taxonomic groups are included in only one or very few publications and clearly lack recurrent evidence for or against monophyly. In order to avoid unduly inflating the number of meaningful tests to be conducted, we therefore used the informative publication count as an adaptive filter. We tested candidate minimum publication thresholds ranging from two to 20 publications, applied Benjamini-Hochberg correction (Benjamini and Hochberg 1995) to the groups passing the filter, and selected the lowest threshold that maximized the number of discoveries at q ≤ 0.05. This procedure selected a minimum of five publications using the TimeTree database, leaving 5,395 groups in the final testing family. Because the cutoff was selected from the observed rejection counts, we included the complete threshold-selection procedure in the end-to-end null simulations described below.

At q ≤ 0.05, 516 NCBI taxa met the threshold for statistically supported recurrent contradiction of monophyly and were included in the catalog of flagged taxa (1.96% of all represented groups, Figs. 1B and 2). None of the 100 end-to-end global-null simulations selected any group. In held-out prediction, partial pooling improved the publication-specific background estimate relative to a rank-wide rate, and adjustment for focal-group sampling improved it further (Fig. 2C). The sampling-detection model likewise outperformed an overall-rate model and a focal-member-count-only model when complete publications or complete broad taxonomic regions were held out during fitting (Fig. 2D, see Supplementary Methods for details).

**Figure 2.**
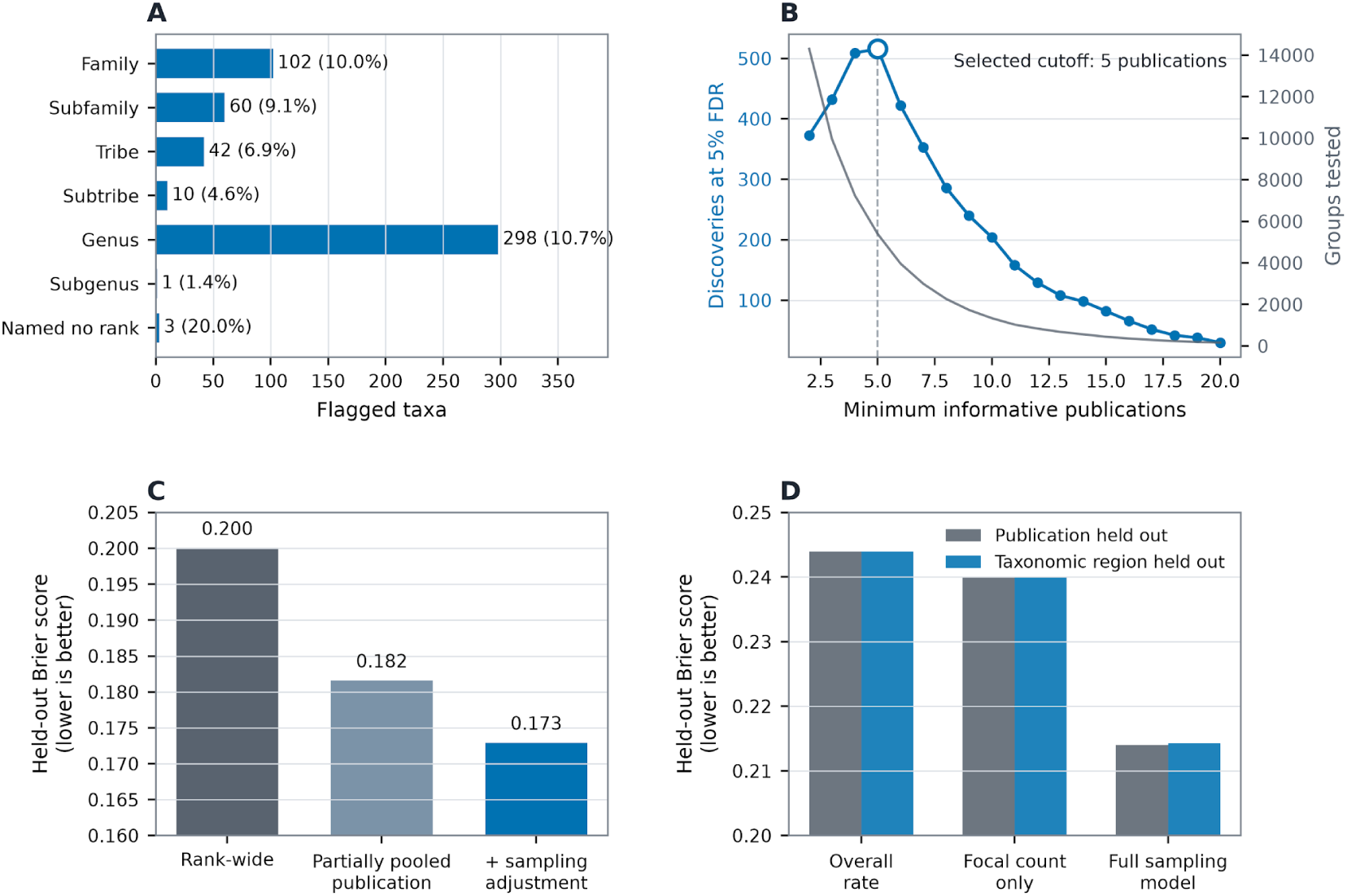
Discovery of NCBI groups reproducibly contradicted by published phylogenies. **(A)** Number and percentage of filtered groups selected at each taxonomic rank, shown in rank order. **(B)** Number of discoveries (blue) and groups tested (gray) as the minimum informative-publication count changes. Five publications produced the largest number of Benjamini-Hochberg-adjusted discoveries. The lowest cutoff was used when counts tied. **(C)** Held-out Brier scores for a rank-wide conflict rate, a partially pooled publication-specific estimate, and the final estimate after adjustment for focal-group sampling. **(D)** Held-out Brier scores for sampling-detection models of increasing detail when complete publications or complete broad taxonomic regions were withheld during fitting. Lower Brier scores indicate better prediction.

Several flagged taxa correspond to groups whose membership has already been challenged or revised in the taxonomic literature. Examples include the placement of *Aeluroscalabotes* in NCBI’s Gekkonidae, the interleaving of *Acanthurus* and *Ctenochaetus*, and legacy generic circumscriptions of *Thelypteris* and *Trichomanes* that predate phylogenetically based revisions (Jonniaux and Kumazawa 2008; Gamble et al. 2011; Sorenson et al. 2013; Ebihara et al. 2006; Almeida et al. 2016; Fawcett et al. 2021). For a subset with adequate species overlap across studies, we separately asked whether the interleaved relationships themselves recurred. Each pair of studies was pruned to the same species, relationships entirely within either taxonomic group were collapsed, and the remaining unrooted cross-group topology was compared using mutual clustering information (Smith 2020). Species labels were then shuffled within their NCBI groups to obtain the null distribution. All 132 cases that had been selected without using this topology statistic and had sufficient shared-species overlap for the secondary analysis remained significant with 9,999 new permutations, and 121 had unadjusted P ≤ 0.001. Thus, the strongest mixed cases contained reproducible topological information rather than only repeated, inconsistent taxonomic intermixing (see Supplementary Methods for details).

Not all unflagged taxa have equally strong evidence of monophyly. To distinguish a low estimated rate of recurrent contradiction supported by many informative studies from one based on sparse evidence, we calculated upper confidence limits on the model’s recurrence parameter for 18,109 taxa with informative publications (Supplementary Data). These limits summarize how strongly the available study trees constrain the possibility of recurrent contradiction (Supplementary Fig. S6), providing information that complements the list of significantly flagged taxa.

### Validation against phylogenomic reference trees

To test whether the catalog of flagged taxa identifies NCBI taxonomic groups that genuinely account for disagreement with independent phylogenies, we represented the complete NCBI taxonomy and a modified version excluding only the flagged taxa’s monophyly requirements (targeted exclusion) as multifurcating species trees and compared both with 13 phylogenomic reference trees published from 2020 through 2026. Each reference used genome-scale data and either dense species sampling or broad coverage of the focal group’s major lineages (Kuderna et al. 2023; Foley et al. 2023; Smith et al. 2023; Hughes et al. 2023; Stiller et al. 2024; Title et al. 2024; Egan et al. 2024; Melendez-Vazquez et al. 2025; Scarpetta et al. 2026; Gable et al. 2022; Nesi et al. 2021; Morales et al. 2025; McGowen et al. 2020). We assessed disagreement using contradiction-aware normalized Robinson-Foulds distance (c-nRF), which counts splits present in only one tree that are incompatible with at least one split in the other, and divides that count by the total number of resolved splits in the two trees. A compatible relationship left unresolved in either tree is not counted as a contradiction. When a reference publication was present in the TimeTree database, we used the catalog fitted after excluding the entire cross-validation fold containing that publication, so the reference publication did not contribute to the exclusions evaluated against its own tree. Twelve references contained at least one flagged taxon whose monophyly requirement was excluded from the comparison. Relative to the complete NCBI taxonomy, the modified version lowered c-nRF for 11 references and raised it slightly for one (mean reduction 0.0436, 95% bootstrap interval 0.0250-0.0649, exact sign-flip *P* = 0.000488). Pooled c-nRF fell from 0.1640 to 0.1235 (Fig. 3A, see Methods for details).

**Figure 3.**
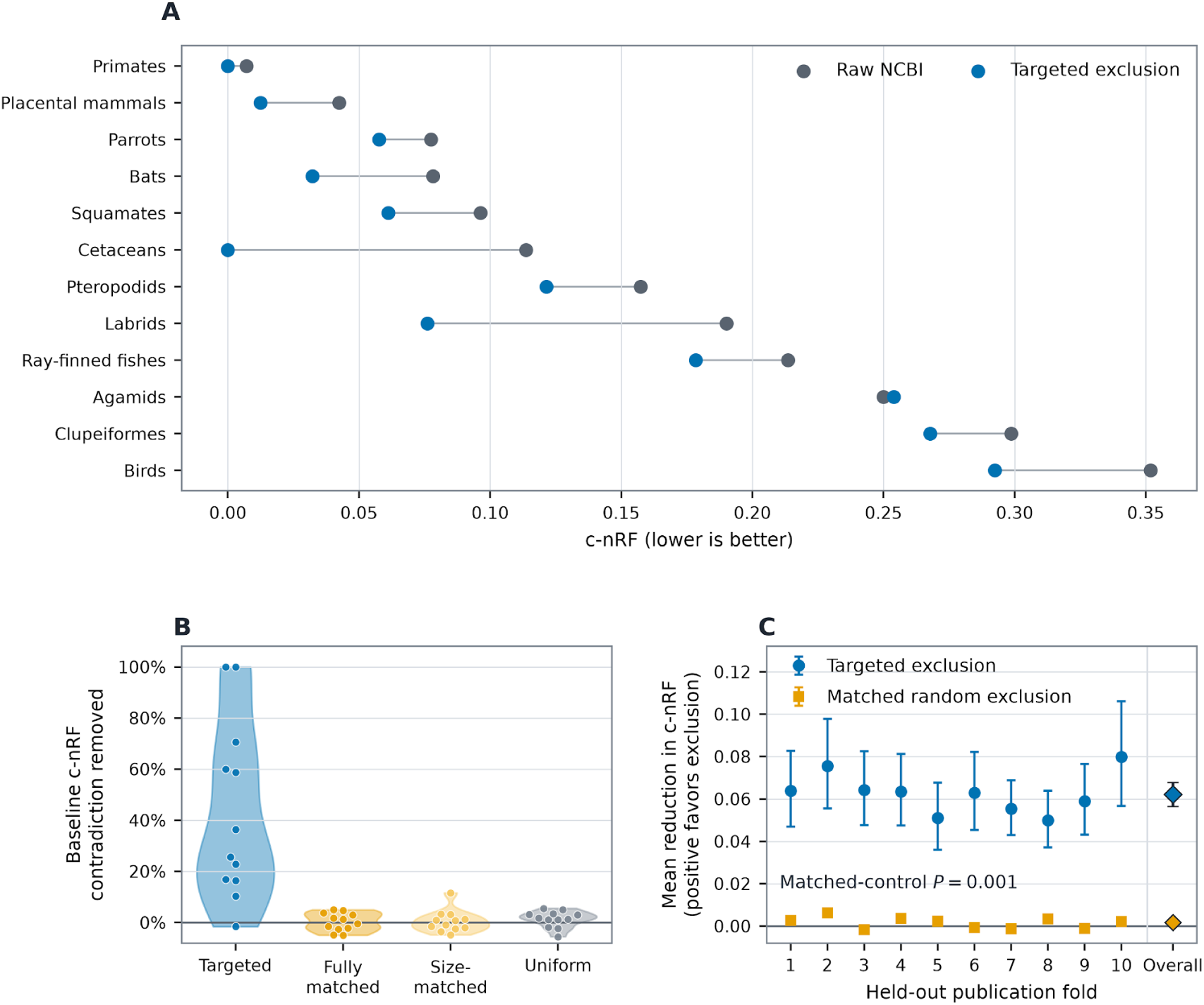
Concordance with published phylogenies. **(A)** c-nRF for complete NCBI taxonomy and targeted exclusion in the 12 affected phylogenomic reference studies. When a reference publication occurred in the discovery collection, its catalog of flagged taxa was fitted after holding out the entire cross-validation fold containing that publication. Lines connect results evaluated against the same reference. **(B)** Distributions across the 12 affected references of the fraction of baseline c-nRF removed by targeted exclusion or random exclusion. Points represent individual references. For each random control, the value is the mean of 10,000 sham catalogs matched by rank, depth, size, and study coverage; by size alone; or sampled uniformly. **(C)** Publication-blocked tenfold cross-validation across the TimeTree study collection. All trees from a publication were assigned to the same fold. Taxa were flagged using the other nine folds, and their monophyly requirements were excluded only when evaluating the withheld fold. Blue circles show mean improvement in c-nRF for targeted exclusion versus the raw NCBI taxonomy for each fold. Orange squares show corresponding results for 1,000 matched random exclusion controls. Diamonds show the overall means. Error bars are 95% intervals obtained by resampling publications. Positive values indicate less contradiction after targeted exclusion.

Because excluding arbitrary monophyly requirements can itself reduce opportunities for contradiction, we generated 10,000 sham catalogs for each of three controls. Each flagged NCBI taxon was replaced by an unflagged taxon after matching taxonomic rank, depth in the NCBI hierarchy, represented clade size, and informative-publication count; after matching represented clade size alone; or by uniform sampling. Every sham catalog was converted into a modified taxonomy by the same procedure used for the catalog of flagged taxa. Their mean c-nRF improvement was 0.0436 for the catalog of flagged taxa compared with 0.0010, 0.0009, and 0.0028 for the three controls, respectively. None of 10,000 replicates equaled or exceeded the observed targeted exclusion result (one-count empirical *P* = 0.0001, Fig. 3B).

We next asked whether a catalog of flagged taxa selected from one subset of publications predicted conflicts in publications across the TimeTree study set that had not contributed to their selection. Of the 5,865 TimeTree phylogenies, 4,555 contained at least four uniquely mapped species and could therefore be compared topologically. We used publication-blocked tenfold cross-validation so that no publication was evaluated with exclusions based on taxa it had helped flag. All trees from each publication were assigned to the same fold. For each fold, background estimation, recurrence testing, publication-count filtering, and false-discovery-rate correction were repeated using only the other nine folds, and the resulting exclusions were evaluated only against trees in the withheld publications (see Methods for details). The training-only catalogs changed at least one represented NCBI split in 858 trees from 825 withheld publications. Relative to complete NCBI, targeted exclusion reduced c-nRF in 654 publications, produced no change in 103, and increased c-nRF in 68. Mean publication-level c-nRF decreased from 0.1882 to 0.1194 (mean reduction 0.0689, 95% bootstrap interval 0.0624-0.0756, one-sided sign-flip *P* = 10^−5^, Fig. 3C). All ten folds had a positive mean effect (range 0.0511-0.0804, one-sided sign test *P* = 0.00098), showing that conflicts learned from one set of publications predicted taxonomic contradictions in publications excluded from selection. We also applied a matched-random control to the held-out-fold evaluation. A distinct same-size control could be constructed for 826 of the 858 affected trees, representing 794 publications. On this common set, targeted exclusion reduced mean publication-level c-nRF by 0.0621, whereas 1,000 matched-random exclusion replicates produced a mean reduction of 0.00161 (95% randomization interval 0.00129-0.00194, empirical *P* = 0.000999, Fig. 3C). Thus, the held-out improvement was not explained simply by representing fewer taxonomic clades.

### Validation against the publication record

To determine whether the taxa flagged by TaxaAudit were known to be unreliable in the publication record, we performed a literature review. We established two cohorts, one consisting of the 516 taxa flagged by TaxaAudit for recurrent contradiction of monophyly, and one consisting of an equal number of taxa with zero conflicting topologies in the TimeTree database. In both cohorts, the number of informative studies per taxon ranged from 5-37 with a median of 8 to minimize any effect of imbalanced support in our assessment. We searched OpenAlex (Priem et al. 2022) for publications that directly challenged the monophyly of each taxon in the two cohorts, either explicitly in the text or through a contradictory inferred topology. We also reviewed a third cohort of 516 otherwise randomly selected taxa with the same range and median informative study count (individual counts and publication references for all three cohorts are in Supplementary Dataset 1).

After excluding publications that contributed trees to the discovery collection, we found qualifying challenges for 231 of 516 taxa flagged by TaxaAudit (44.8%), compared with 64 of 516 controls (12.4%), a 3.61-fold difference (Fig. 4).

**Figure 4.**
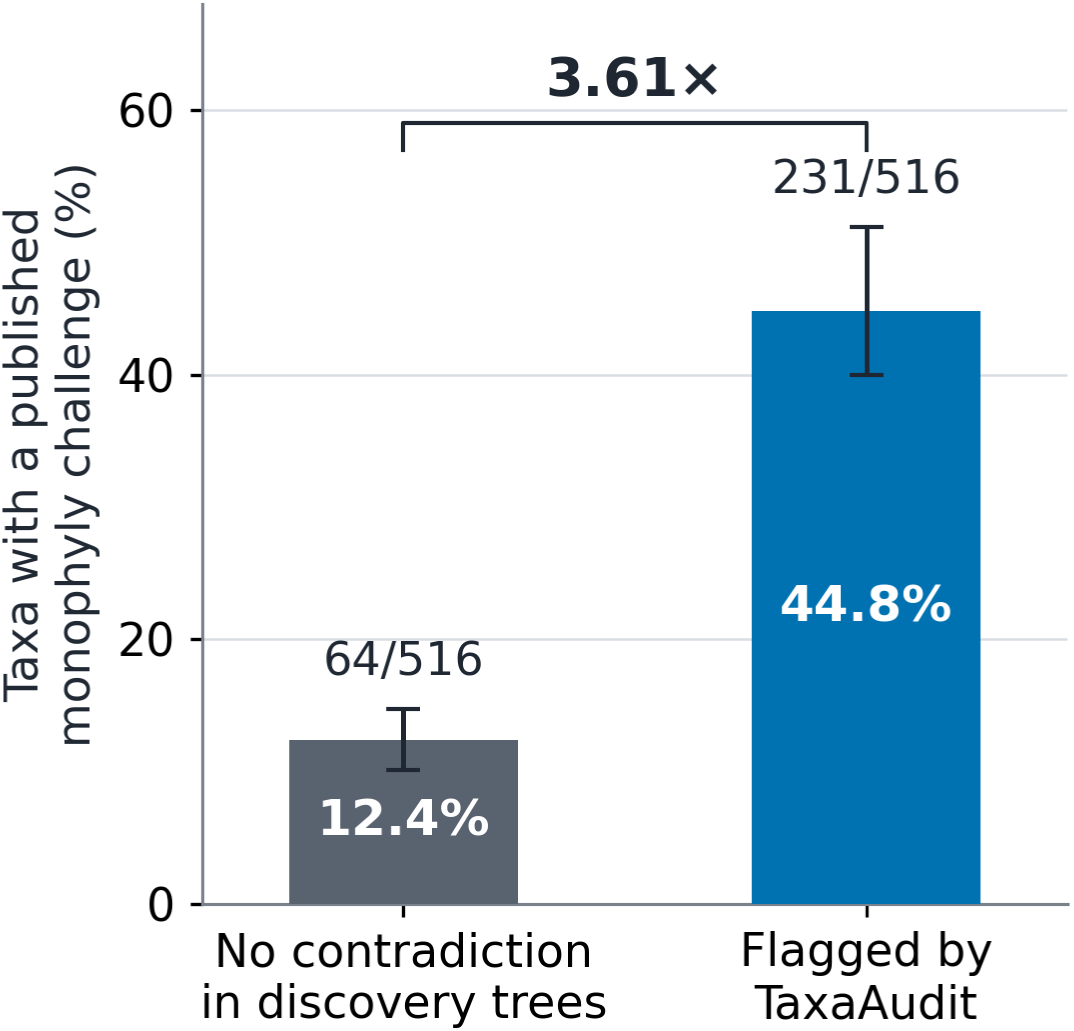
Published monophyly challenges are enriched among taxa flagged by TaxaAudit. Results are shown for two cohorts of 516 taxa, one consisting of taxa flagged by TaxaAudit, and one consisting of taxa with no conflicting topologies in the TimeTree study set. Both cohorts had a median of 8 informative studies per taxon and a range of 5-37 informative studies. Labels give percentages and numbers of taxa with a report divided by cohort size. Error bars are 95% confidence intervals from 20,000 bootstrap resamples of taxonomic orders, with class or phylum fallback.

### Tree-inference experiments

Finally, we tested whether the complete NCBI hierarchy or the modified version with flagged taxa excluded from monophyly constraints provided more accurate guidance for maximum-likelihood tree inference. We inferred maximum-likelihood trees under three treatments. The first required all represented NCBI groups to be monophyletic. The second excluded the monophyly requirements of flagged taxa, using the catalog fitted after excluding the reference publication’s cross-validation fold. The third used no constraint. A set of 16 aligned matrices was assembled from public GenBank records as described in Methods (Allard et al. 2026). These averaged 72,667 aligned sites. We repeated inference after downsampling to approximately 7,735 sites and 1,500 sites (see Methods, Supplementary Fig. S4). The targeted-exclusion constraint reduced c-nRF relative to the complete NCBI constraint when the full data were used (mean improvement 0.04285, 95% publication-bootstrap interval 0.01324-0.07724, exact *P* = 0.0146), at the 7,735-site target (0.03705, interval 0.01204-0.06687, *P* = 0.0117), and at the 1,500-site target (0.03361, interval 0.00799-0.06599, *P* = 0.0254, Fig. 5A). Publication-level effects are shown in Supplementary Fig. S1.

**Figure 5.**
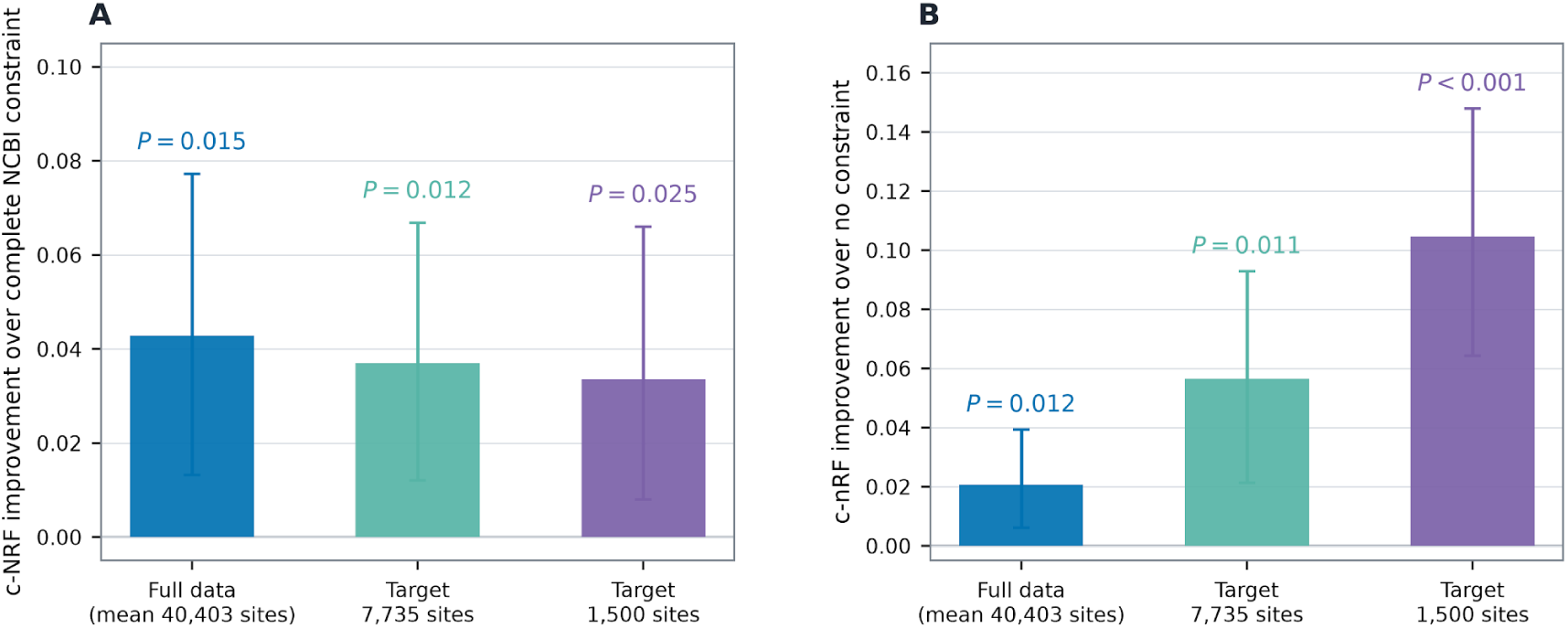
Excluding flagged taxa from monophyly constraints improves maximum-likelihood inference while retaining useful taxonomic guidance. Both panels show the same full-data and reduced-data treatments. Positive values favor targeted exclusion. **(A)** Improvement in c-nRF relative to the complete NCBI constraint when inferred trees were compared with held-out phylogenomic reference studies. **(B)** Improvement relative to no constraint after excluding reference relationships that conflict with complete NCBI guidance or are explicitly imposed by the targeted-exclusion constraint. Panel B therefore measures recovery of reference relationships that all treatments could recover but that targeted exclusion did not hard-code. Bars show means giving each publication equal weight. Error bars are 95% bootstrap intervals from 100,000 samples in which the 14 publications were sampled with replacement. Labels give two-sided exact tests obtained by reversing the sign of each publication’s complete contribution.

We next asked whether the taxonomic groups retained after targeted exclusion improved inference beyond relationships directly imposed by the constraint. From each held-out phylogenomic reference tree, we excluded every split incompatible with the complete NCBI constraint and then excluded every remaining split explicitly present in the targeted-exclusion constraint. All scored relationships were therefore possible under complete NCBI guidance, but none was hard-coded by the targeted constraint. Relative to unconstrained inference, targeted exclusion reduced c-nRF for this partial reference topology with full data (mean improvement 0.02076, 95% publication-bootstrap interval 0.00611-0.03944, exact *P* = 0.0117), at the 7,735-site target (0.05655, interval 0.02131-0.09301, *P* = 0.0107), and at the 1,500-site target (0.10465, interval 0.06437-0.14803, *P* = 0.00037, Fig. 5B, see Methods for details). Thus, the benefit extended to relationships inferred from the sequence matrix rather than only those enforced by retained taxonomic groups.

The less favorable comparison against complete reference trees was explained by taxonomic disagreement that targeted exclusion had not removed. Across publications, the targeted-exclusion advantage over no constraint became smaller as the fraction of reference splits incompatible with complete NCBI increased (Spearman *ρ* = −0.722, *P* = 0.00486 with full data; *ρ* = −0.723, *P* = 0.00458 at the 7,735-site target; and *ρ* = −0.682, *P* = 0.00945 at the 1,500-site target, Supplementary Fig. S2). This association disappeared when incompatible reference splits were excluded. A baleen-whale example is shown in Supplementary Fig. S3.

BUSCO lineage datasets illustrate a widely used application of NCBI-derived taxonomic groups: defining sets of conserved single-copy orthologs for assessing genome completeness and constructing phylogenies (Tegenfeldt et al. 2025). Synechococcaceae, a family that names one such dataset, was contradicted in all six informative publications in our discovery collection (q = 0.0482). We examined additional published phylogenomic trees to assess this example. In two 650-genome analyses based on 834 markers and 31 core genes, 74 mapped genomes assigned to this family by NCBI did not form an exclusive clade. Marine members grouped with Prochlorococcus, whereas JA-3-3Ab and PCC 7336 occupied a distant lineage. A branch demonstrating this separation had reported support of 100 in both analyses (Chen et al. 2021). A separate 260-locus phylogeny showed corresponding separations and supported revised taxonomic assignments (Komárek et al. 2020). These results corroborate the TaxaAudit finding and motivate examination of lineage membership and marker applicability, although effects on BUSCO scores were not tested.

## Methods

### Study-tree classification and publication-level observations

Each group was evaluated in the complete rooted study tree. For a sampled NCBI group *G*, monophyly was recovered when its most recent common ancestor contained exactly the sampled members of *G*. A resolved descendant clade containing some, but not all, members of *G* together with nonmembers counted as contradiction. Absence of both was compatible but unresolved and was omitted. The publication was the replication unit. If a publication contained several tree files, the predominant informative status supplied one binary observation. Ties supplied no observation (Fig. 1C, see Supplementary Methods for details).

### Publication-specific background conflict

For publication *i* and group *g*, the null conflict probability was first estimated from all other groups of the same rank in publication *i*. Sparse publication-specific estimates were partially pooled toward the corresponding rate in the full study collection using a beta-binomial empirical-Bayes model (Morris 1983, see Supplementary Methods for details). We then fitted a monotone model using that estimate and the number of sampled members of *g*. Five-fold cross-fitting kept every observation of the focal NCBI group out of the model that predicted its background probability. This estimates how often a comparable apparent conflict would be expected from the publication and its sampling even if group *g* had no recurring problem (see Supplementary Methods for details). The intuition for this approach is that a study that used very little data may be expected to have an elevated level of phylogenetic noise leading to mixing of species even between true clades.

### Sampling-aware test of recurrent non-monophyly

Sampling also determines whether a real problem can be observed. We trained the sampling-detection model on 82,000 observations, comprising 2,000 complete-tree examples and 80,000 reduced-tree observations. The complete examples were trees in which a focal NCBI group was contradicted. Each reduction independently sampled focal-group members, nonmembers under the focal group’s immediate NCBI parent, and more distant nonmembers, after which the same complete-tree classifier was rerun. Represented sibling lineages were counted after sampling. A monotone gradient-boosted tree model predicted whether the reduced tree retained the original contradiction from the numbers of focal members, immediate-parent nonmembers, and represented sibling lineages. Complete publications or complete orders or higher-ranking taxa were held out during predictive validation, and production predictions were cross-fitted by broad taxonomic region (Figs. 2D and S5, see Supplementary Methods for details). The fitted probability model had two mutually exclusive routes to observed conflict: ordinary background discordance, or a recurring group-specific violation that the publication’s sampling exposed. The only fitted parameter for a focal group was the strength of the latter recurrence. We compared the best-fitting value with the null value of zero. For groups passing a conservative asymptotic screen, 99,999 parametric-bootstrap data sets were simulated from their publication-specific null probabilities, and the recurrence parameter was refitted to each. The likelihood-ratio statistic compared the best-fitting model with the null model. The group *P* value was the proportion of null replicates with an equal or larger statistic, including the one-count correction (see Supplementary Methods for details).

### Publication-count filtering and multiple-testing correction

Informative publication count was used to exclude groups with too little evidence to test recurrence effectively before multiple-testing correction. We evaluated minimum counts from two through 20, applied Benjamini-Hochberg correction within each retained set, selected the lowest minimum attaining the largest number of discoveries, and reported *q* ≤ 0.05 (Benjamini and Hochberg 1995). The complete adaptive rule, its relationship to independent hypothesis filtering, and its end-to-end null calibration are described in the Supplementary Methods.

### Validation against published phylogenies

We represented the complete NCBI taxonomy and the targeted-exclusion version as multifurcating species trees. Targeted exclusion omitted a flagged taxon’s exact monophyly requirement while retaining all species and the monophyly requirements of unflagged ancestors and descendants. Comparisons used c-nRF, calculated from incompatible unrooted splits. Reference publications present in the discovery collection were evaluated using catalogs fitted after excluding their entire cross-validation fold. For the study-wide comparison, all trees from a publication remained in the same fold, and its outcomes were excluded from background fitting and selection of flagged taxa. The sampling-detection model was held fixed across these catalog fits.

Random-exclusion controls tested whether excluding comparable monophyly requirements could explain the improvement. Reference-tree controls replaced targeted taxa with nontarget taxa under full matching, size-only matching, or uniform sampling. Cross-validation controls removed the same number of represented taxonomic relationships from each held-out comparison, using matched nontarget groups. Effects were combined within publications before statistical testing. Control selection, c-nRF normalization, and publication-level resampling are detailed in the Supplementary Methods.

### Literature review of published monophyly challenges

We compared the 516 flagged taxa with 516 controls matched approximately for informative-publication count and having no observed contradiction in the discovery trees. Candidate publications retrieved through OpenAlex underwent agent-assisted review of available abstracts and selected full texts (Priem et al. 2022). A qualifying report explicitly challenged the focal taxon’s monophyly, presented an incompatible topology, or reassessed its circumscription on that basis. We excluded reports from discovery publications and reports with unresolved publication identities. Unassessable papers were excluded individually, retaining all taxa in both cohorts. We compared the fractions with at least one remaining report using taxonomic-region bootstrap intervals and assessed sensitivity to rank and publication coverage. Full eligibility rules, publication matching, statistics, and the descriptive third cohort are documented in the Supplementary Methods and Dataset 1.

### Maximum-likelihood inference experiments

Molecular matrices were assembled from GenBank sequences through homology clustering, sequence and locus quality control, alignment, and trimming (Allard et al. 2026). We restricted matrices to species shared with each phylogenomic reference and analyzed full data and two reduced levels targeting 7,735 or 1,500 sites. IQ-TREE analyses differed only in constraint treatment: complete NCBI, targeted exclusion, or no constraint (Nguyen et al. 2015; Minh et al. 2020). Targeted constraints used catalogs excluding the reference publication’s cross-validation fold.

We scored inferred trees against complete references and, separately, against reference relationships compatible with complete NCBI but not explicitly imposed by the targeted constraint. The latter comparison assessed recovery of relationships that all treatments could infer from the data. Replicates were averaged within cases, and cases were combined within publication using the number of scored reference splits. Each publication contributed equally to the overall mean and statistical tests. The Supplementary Methods specify matrix assembly, coverage-preserving reductions, inference settings, scoring, and publication-level uncertainty estimation.

## Discussion

The NCBI taxonomy offers an accessible evolutionary framework for research that compares species, but this use depends on the implicit assumption that named groups correspond to evolutionary clades. TaxaAudit evaluates that assumption using the accumulated phylogenetic literature. We identified 516 NCBI groups whose monophyly is repeatedly contradicted. An independent literature review found substantially more published monophyly challenges among flagged taxa than among controls. Excluding flagged taxa from monophyly requirements also improved agreement with independent phylogenomic reference trees and publications withheld from discovery. These findings support our ability to identify unreliable clade assumptions systematically rather than relying on detailed familiarity with the phylogenetic literature for each taxon.

The flagged taxa should not automatically be interpreted as errors in the NCBI taxonomy. As the whale example illustrates, a classification retained in NCBI can conflict with published molecular phylogenies (Árnason et al. 2018; McGowen et al. 2020). In other cases, the taxonomy remains highly disputed, and curators may reasonably take a conservative approach to contentious relationships (Schoch et al. 2020). TaxaAudit does not adjudicate these competing classifications. Instead, it identifies groups for which assuming monophyly is difficult to reconcile with the sampled phylogenetic literature. Its value lies in making those disagreements visible to researchers who might otherwise be unaware of them.

These disagreements matter whenever taxonomic membership determines which organisms or reference data are treated as a coherent evolutionary group. For example, researchers may select BUSCO lineage datasets using NCBI taxonomic assignments, relying on them to identify appropriate sets of conserved single-copy orthologs for genome assessment or phylogenetic analysis (Tegenfeldt et al. 2025). Our finding for *Synechococcaceae* illustrates why that choice may warrant scrutiny: a marker set derived from some members of a non-monophyletic family may not adequately represent other species assigned to it.

Identifying recurrent contradictions is only one use of the model. The upper confidence limits on recurrence provide complementary information for groups not flagged as problematic. A group with no observed contradiction may be consistent with the published phylogenies or merely poorly sampled. By distinguishing those cases, TaxaAudit can help researchers select groups with stronger published evidence for monophyly before undertaking analyses that depend on it.

Flagged taxa also provide candidates for reassessment by taxonomic specialists. The catalog directs attention toward cases in which published molecular phylogenies repeatedly challenge the membership of named groups, without prescribing a particular reclassification. Such prioritization could help focus taxonomic research in response to the biodiversity crisis (Savage 1995; Singh 2002).

The approach is also highly flexible in terms of the taxonomy being interrogated and the reference topology set used for comparison. For example, we tested NCBI taxa at family rank and below, but we provide exploratory results for higher ranks through phylum (Supplementary Fig. S7). The same framework could also be applied using tree collections tailored to a particular research question, including morphological phylogenies, timed or otherwise, and curated samples of trees inferred from different genomic datasets or using competing inference methods. This allows the active systematist to carry out rapid hypothesis testing and ensure the robustness of their proposed taxonomic actions to different data types or methods.

The present analysis treats study trees as evidence without assigning additional weights based on publication metadata. Our background-conflict and sampling models may already capture some differences in the informativeness of individual studies, but researchers may also wish to weight publications by molecular data quantity, recency, or other criteria. The model permits such weights, although we did not use them here, and their effects require further evaluation.

More generally, TaxaAudit makes information from the phylogenetic literature accessible for decisions about evolutionary groupings. This information is relevant when researchers select species for comparative analyses, reconstruct gene-family histories, assemble ortholog sets, or synthesize phylogenies (Smith et al. 2009; Hinchliff et al. 2015; Kumar et al. 2022; Kuznetsov et al. 2023; Barrera-Redondo et al. 2023). By helping researchers identify groups with recurrent evidence against monophyly and assess the strength of evidence for other groups, TaxaAudit allows the NCBI taxonomy to serve not merely as a system for organizing biological data, but as an evolutionary framework whose clade assumptions can be evaluated against published evidence.

## Acknowledgments

This work was supported by a research grant from the National Institutes of Health to S.K. (R35GM139540).

## Data and code availability

The TaxaAudit source code, catalog of flagged taxa, and data on upper confidence limits for recurrent contradiction are available at https://github.com/John-Allard/TaxaAudit. The study trees are available from TimeTree (https://timetree.org/). Additional supplementary datasets will be made available.

## Notes

https://github.com/John-Allard/TaxaAudit

### Competing Interest Statement

The authors have declared no competing interest.

