## Supplementary methods and figures for "TaxaAudit: A statistical model of consensus in phylogenetics identifies non-monophyletic taxa"

### 1 Supplementary Methods

#### 1.1 Study trees and taxonomic mapping

##### 1.1.1 Input collection and taxonomic mapping

The input comprised 5,865 molecular study-tree files from the TimeTree literature collection. TimeTree records published molecular divergence-time analyses and their source trees (Hedges et al., 2006; Kumar et al., 2017, 2022). The publication, identified by PMID when available and otherwise by its TimeTree citation key, was the unit of replication. When several trees from a publication contained the same group, their informative classifications were combined into one observation for that publication and group.

The catalog of flagged taxa contains taxa with statistically supported recurrent contradiction of monophyly after the publication-count filter and multiple-testing correction described below.

Taxonomic membership was evaluated with the NCBI Taxonomy downloaded 22 July 2026; the downloaded archive had SHA-256 `9706198c129c6b70ec591009431f742649b939d5de86e102d469e6928034a1fe`. Retired identifiers were followed through `merged.dmp`; identifiers appearing only in `delnodes.dmp` were not used. Named unranked clades were treated in the same way as named ranked groups. Administrative or placeholder nodes denoting unclassified records, environmental samples, or *incertae sedis* placements were not tested as biological groups, but their named descendant groups remained eligible.

Tree tips contributed only when their candidate identifiers resolved to exactly one NCBI species and that species appeared once in the tree. Of 515,804 input tips, 477,749 had a single species-level candidate. These represented 468,255 unique species occurrences. Another 9,494 tip occurrences were excluded because two or more tips in the same tree mapped to the same species, 17,912 had only non-species candidates, and 20,143 had no taxonomic candidate. The complete mapping attrition is supplied with the data release.

##### 1.1.2 Groups eligible for observation

The screened ranks were family, subfamily, tribe, subtribe, genus, subgenus, species group, and species subgroup. Named "no rank" clades below family were also included; the lack of a Linnaean rank does not make a named clade uninformative. Orders, classes, and other higher ranks were outside the predefined scan. A tree could inform group $g$ when it contained at least two mapped members of $g$ and at least one mapped species outside $g$. One sampled member is necessarily monophyletic and cannot reveal failure of the group. A group represented in fewer than two publications could not show recurrence, but such groups were retained in the initial universe so that the later publication-count filtering step had an explicit denominator. The resulting universe contained 26,291 represented NCBI taxa.

#### 1.2 Classifying a group's status in a study tree

Let $G$ be the sampled members of an NCBI group and let $U$ be every uniquely mapped species in the same complete tree. We tested whether $G$ formed a clade in the rooted study tree. We assigned one of three states:

1. **Displayed**: the most recent common ancestor of $G$ had exactly the members of $G$ among its mapped descendants.
2. **Contradicted**: at least one resolved child subtree below that ancestor contained some, but not all, sampled members of $G$ together with at least one nonmember. This test uses ancestor-descendant relationships in the rooted tree.
3. **Compatible but unresolved**: the tree did not display $G$, but no resolved split contradicted it. A refinement of the polytomy could therefore display $G$.

The third state was treated as missing information. This distinction prevents a consensus-tree polytomy from being scored as non-monophyly merely because it does not display the proposed clade. The classification yielded 76,360 assessments of groups within individual trees. The later c-nRF comparisons used unrooted splits, as defined in the validation section.

For group $g$ in publication $i$, compatible-but-unresolved tree files were first omitted. If the remaining files had a unique modal state, displayed or contradicted, that state supplied one publication-level binary observation $Y_{ig}$; a tie was omitted. Thus $Y_{ig} = 1$ denoted a resolved contradiction and $Y_{ig} = 0$ denoted displayed monophyly. Of the 26,291 represented taxa, 18,109 had at least one informative publication. These taxa contributed 73,181 informative assessments of individual taxa within publications across 3,392 publications. The remaining 8,182 represented taxa had no informative publication-level outcome. Publications outside those 3,392 supplied no informative observation for any eligible group.

#### 1.3 Sampling-aware recurrence model

##### 1.3.1 Overview

The number or fraction of contradicting publications is not comparable across groups. Publications differ in their general rate of apparent taxonomic conflict, and a sparsely sampled group has less opportunity to reveal a violation. We therefore modeled

$$\Pr(Y_{ig} = 1 \mid \rho_g) = f_{ig} + (1 - f_{ig}) D_{ig} \rho_g. \tag{1}$$

Here $f_{ig}$ is the probability of ordinary apparent conflict expected for a comparable group in publication $i$; $D_{ig}$ is the probability that publication $i$ would expose a real violation under its sampling; and $\rho_g \in [0, 1]$ is the excess recurring tendency specific to group $g$. Under the null hypothesis $\rho_g = 0$, the conflict probability is $f_{ig}$. A displayed clade in a sparsely sampled study supplies less counterevidence because it has a lower $D_{ig}$, while a resolved contradiction is still an observed contradiction.

##### 1.3.2 Empirical-Bayes estimate of ordinary conflict

We first estimated the ordinary conflict rate from other groups of the same NCBI rank in the same publication. For a focal observation, let $x_{ig}$ be the number of contradictions and $n_{ig}$ the number of informative outcomes after removing the focal group from that publication-rank cell. Let $r_{ig}$ be the rate among all observations of that rank after removing every publication-level observation of focal group $g$:

$$r_{ig} = \frac{X_{-g} + 0.5}{N_{-g} + 1}. \tag{2}$$

The initial background estimate was

$$f_{ig}^{(0)} = \frac{x_{ig} + s r_{ig}}{n_{ig} + s}. \tag{3}$$

Equation 3 is the posterior mean under a binomial likelihood and a beta prior with mean $r_{ig}$ and concentration $s$, making it a beta-binomial empirical-Bayes model (Morris, 1983). A publication with many same-rank observations is represented mainly by its own rate; a publication with few observations is pulled toward the collection-wide rate for that rank. The fitted $s = 3.3724$ is the effective number of observations supplied by the broader rate.

The shrinkage strength was selected by leave-one-observation-out predictive likelihood (Stone, 1974). For every candidate $s$, each binary observation was temporarily omitted, equation 3 was recalculated without it, and the log probability assigned to its observed value was recorded. We selected the $s$ maximizing the sum of those held-out log probabilities. This procedure chooses how strongly sparse publication estimates should be pulled toward the broader rank average; it does not predict phylogenetic topology.

Group sampling also changes the chance of ordinary apparent conflict. We therefore fitted a gradient-boosted decision-tree classifier (Friedman, 2001), using the histogram implementation in scikit-learn (Pedregosa et al., 2011), with two inputs: the logit of $f_{ig}^{(0)}$ and $\log(1 + m_{ig})$, where $m_{ig}$ is the number of sampled members of $g$. The fitted probability was the final $f_{ig}$. Five-fold group cross-validation assigned all observations of one NCBI group to the same fold, so the model predicting $g$ had been trained without any outcome from $g$. Both effects were constrained to be nondecreasing. The model used learning rate 0.05, 200 iterations, at most 15 leaves, at least 100 observations per leaf, and L2 regularization 2.0. The rank-wide conflict rate, the partially pooled publication estimate, and the final sampling-adjusted background had cross-fitted Brier scores of 0.1999, 0.1816, and 0.1729, respectively (Main Fig. 2C); lower scores indicate better prediction. The final model's mean log loss was 0.5202. A sensitivity model also included the number of sampled nonmembers, but this over-corrected groups in large trees because most distant nonmembers cannot plausibly invade the focal clade; the prespecified primary model used focal-member sampling only.

##### 1.3.3 Probability that sampling would expose a violation

The detection term $D_{ig}$ was learned by reducing the species samples of complete study trees in which a focal NCBI group was contradicted. We selected 2,000 contradicted tree-group observations, stratified across taxonomic ranks and tree-size classes. They came from 686 publications and 195 broad taxonomic regions. For each template, we retained the complete observation and generated 40 reduced observations, giving 82,000 training records: 2,000 complete and 80,000 reduced records. Each reduction independently sampled the focal-group members, nonmembers under the focal group's immediate NCBI parent, and more distant nonmembers. The number of represented sibling lineages under the parent was counted after sampling. We then reran the same three-state complete-tree classifier used in the primary scan. The binary response was one when the reduced tree still contradicted the focal group and zero otherwise. Thus, the training target directly measured whether a particular species sample retained an already observed complete-tree contradiction. Each original template received equal total weight, regardless of how many valid reductions it produced.

Templates were allocated across strata defined by NCBI rank and the integer part of $\log_2$ of the number of mapped species in the tree, initially allocating an equal number to each stratum and filling any remaining places from unselected observations. Template selection and reductions used seed 20260820, with independent streams derived from the tree-group identifier and draw number. For each of the three species pools, let $n$ be its available count and $m$ the minimum retained count (two for focal members and zero for each nonmember pool). If $n \leq m$, all were retained. Otherwise, all were retained with probability 0.12. In the remaining draws, a retention fraction was sampled uniformly on the log scale from $\max(m/n, 0.01)$ to one, followed by a binomial count with parameters $n$ and that fraction, bounded below by $m$. Species were sampled uniformly without replacement within each pool. If neither nonmember pool retained a species, one was restored from the local pool when available, or otherwise from the distant pool.

We compared monotone logistic and monotone histogram gradient-boosting classifiers (Friedman, 2001; Pedregosa et al., 2011). Predictors considered the number of sampled focal-group members, sampled nonmembers under the focal group's immediate NCBI parent, sampled nonmembers outside that parent, and the number of represented sibling lineages under the parent. A sibling lineage was an immediate daughter of the focal group's parent represented by at least one sampled local nonmember. Counts were transformed as $\log(1+x)$, and every effect was constrained to be nondecreasing. The selected model used focal-group members, immediate-parent nonmembers, and represented sibling lineages. It used learning rate 0.05, 150 iterations, at most 15 leaves, at least 40 observations per leaf, and L2 regularization 1.0.

Predictive validation blocked complete publications or complete broad taxonomic regions into ten folds using scikit-learn's GroupKFold. A region was the focal group's ancestral order, or its class or phylum when the preceding rank was absent; the highest ancestor below the root was used if none of these ranks was available. In each analysis, all records derived from one template remained in the same fold, and the model predicting a fold was trained on the other nine. On the 80,000 reduced observations, with equal total weight per template, the selected model had Brier scores of 0.2140 when publications were held out and 0.2143 when broad taxonomic regions were held out. Corresponding scores were 0.2439 and 0.2439 for a constant-probability model and 0.2399 and 0.2399 for a monotone model using only the focal-member count (Main Fig. 2D). Calibration summaries and alternative model results are supplied in the data release.

Production predictions used five taxonomic-region folds specified before downstream catalog evaluation. For a region represented among the training templates, its predictions came from a model fitted without that complete region. Regions absent from the training collection used the model fitted to all templates. Cross-fitted predictions supplied 94.6% of the 73,181 publication-level observations. Production $D_{ig}$ values had mean 0.691, median 0.719, and 1st-99th percentiles 0.230-0.957.

As an ablation, we repeated the complete analysis with every $D_{ig} = 1$, treating every informative publication as certain to expose a real violation. This assumption had a held-out Brier score of 0.4207 under either blocking scheme (Supplementary Fig. S5B). It selected 456 groups, of which 455 were also in the 516-group primary catalog. The learned model detected 61 additional groups and improved prediction in withheld publications relative to the $D = 1$ catalog (mean all-publication c-nRF advantage 0.00109, 95% publication-bootstrap interval 0.00076-0.00148, one-sided sign-flip $P = 10^{-5}$). Against the 13 phylogenomic references, it was better for six, identical for six, and slightly worse for one (exact one-sided sign-flip $P = 0.0156$ across the seven nonzero differences).

##### 1.3.4 Likelihood-ratio statistic and parametric bootstrap

For a fixed group, the Bernoulli log likelihood was

$$\ell_g(\rho) = \sum_i \left[Y_{ig} \log p_{ig}(\rho) + (1 - Y_{ig}) \log\{1 - p_{ig}(\rho)\}\right], \tag{4}$$

where $p_{ig}(\rho)$ is given by equation 1. We evaluated 201 equally spaced values of $\rho$ from zero through one and calculated

$$T_g = 2\left\{\max_{0\le\rho\le1} \ell_g(\rho) - \ell_g(0)\right\}. \tag{5}$$

Because zero is on the boundary of the parameter space, the one-sided $0.5\chi^2_0 + 0.5\chi^2_1$ mixture (Self and Liang, 1987) was used only as a computational screen. Groups with asymptotic BH $q \le 0.30$ received 99,999 parametric-bootstrap replicates. In replicate $b$, every publication outcome was independently simulated as

$$Y^{*(b)}_{ig} \sim \text{Bernoulli}(f_{ig}), \tag{6}$$

and $\rho$ and $T^{*(b)}_g$ were refitted exactly as for the observed data. The final value was

$$P_g = \frac{1 + \#\{T^{*(b)}_g \ge T_g\}}{100{,}000}, \tag{7}$$

using the one-count correction of Phipson and Smyth (2010). Groups outside the computational screen received $P_g = 1$ for multiple-testing purposes. The screen was deliberately much broader than the final rejection set; no selected group depended on the asymptotic approximation.

###### 1.3.5 Publication-count filtering and multiple-testing correction

Thousands of groups occurred in too few publications to supply appreciable evidence of recurrence. We therefore used the number of informative publications as a power-related filtering variable before multiple-testing correction. Filtering on a covariate related to power can increase detection when the filter preserves the null distribution of the subsequent test (Bourgon et al., 2010). We evaluated minimum counts from two through 20. At each count, Benjamini-Hochberg (BH) correction (Benjamini and Hochberg, 1995) was applied to bootstrap $P$ values only for groups passing that count. We selected the lowest count attaining the largest number of BH discoveries. The selected minimum was five publications: 5,395 groups entered the final family and 516 had $q \le 0.05$. The neighboring cutoffs produced 373 findings at two publications, 432 at three, 509 at four, and 422 at six, with progressively fewer thereafter. The complete distribution of bootstrap $P$ values is shown in Supplementary Figure S5A.

Selecting the filter threshold that maximizes the number of discoveries has precedent in genomic workflows, including DESeq2 (Love et al., 2014), but it is an adaptive choice and is not guaranteed by independent-filtering theory alone (Ignatiadis et al., 2016; Kim and Schliekelman, 2016). We therefore included the entire choice - re-estimation of the background model, recurrence testing, evaluation of every candidate publication-count minimum, selection of the minimum, and BH correction - in the end-to-end global-null calibration described below. This makes five publications an empirically calibrated analysis choice, not an asserted biological minimum. BH-adjusted values are reported as $q$ values. The original BH theorem assumes independent true-null tests (Benjamini and Hochberg, 1995); broader positive-dependence results and the conservative Benjamini-Yekutieli modification describe alternatives under dependent testing (Benjamini and Yekutieli, 2001). Because nested NCBI groups scored in the same trees are dependent, we do not claim a distribution-free guarantee of exactly 5% FDR under arbitrary dependence.

The selected groups had a median of eight informative publications (range 5-37). They included 298 genera, 102 families, 60 subfamilies, 42 tribes, 10 subtribes, three named unranked clades, and one subgenus. Groups below the selected filter were designated evidence-limited and retained, not declared monophyletic.

#### 1.4 End-to-end calibration

To assess the complete primary decision rather than only its one-group likelihood ratio, we performed semisynthetic global-null simulations on the observed publication-by-group design. We retained each group's publications, sample sizes, fixed cross-fitted detection probabilities, rank, and fitted null probability, but generated every outcome from its null Bernoulli probability. For every replicate we then re-estimated the empirical-Bayes background, recalculated recurrence tests and bootstrap values, reevaluated and selected the publication-count filter, and reapplied BH correction. None of 100 replicates selected any group. Thus, the empirical probability of one or more false selections was zero under the fitted global-null model. This experiment preserves the observed publication sampling and group-specific fitted null probabilities. It does not reproduce every dependence among nested or overlapping groups scored in the same trees, nor dependence arising from reuse of molecular data among publications. The calibration therefore supports the complete implemented procedure under the fitted null but is not a proof of FDR control under arbitrary dependence.

#### 1.5 Secondary test of recurring interleaved topology

Repeated contradiction can arise from different misplaced species in every study. We therefore performed a separate topology-recurrence test for 132 high-information endpoint cases that had been selected without using this topology statistic and had adequate shared-species overlap. This analysis is supporting evidence about the structure of strong conflicts and is not a gate in the 516-group primary catalog.

For each pair of publications, the representative conflicting trees were pruned to their shared species, requiring at least two shared species from each focal NCBI group. Internal nodes whose descendants all belonged to only one focal group were collapsed. The remaining "mixed skeleton" retains cross-group structure. For study-tree pair $k$, let $I_k$ be the mean clustering entropy of its two mixed skeletons and $M_k$ their mutual clustering information, calculated from the unrooted splits with TreeDist (Smith, 2020). Because no dummy root-anchor leaf was added, this secondary analysis tests recurrence of unrooted cross-group structure rather than root-dependent nesting direction. For publication $i$, comparisons with all partners $j$ were reduced to

$$s_i = \frac{\sum_j M_{ij}}{\sum_j I_{ij}}, \qquad w_i = \max_j I_{ij}. \tag{8}$$

The case statistic was the information-weighted mean

$$S = \frac{\sum_i w_i s_i}{\sum_i w_i}. \tag{9}$$

This gives one score to each publication. More informative overlaps can contribute more, but comparison with many partners cannot multiply a publication's total weight beyond its best-resolved overlap.

The null shuffled species labels within each focal NCBI group in each publication's complete focal tree and only then pruned to the shared species for every partner. One relabeling of a publication was reused across all its partner comparisons in a replicate, preserving their dependence. The shuffle retains tree shape, resolution, sampling counts, and the overall amount of intermixing but destroys recurrence of species-specific cross-group relationships. With $B = 9{,}999$ replicates,

$$P = \frac{1 + \#\{S^{*(b)} \geq S\}}{B+1}. \tag{10}$$

All 132 cases had unadjusted permutation $P \leq 0.05$ and 121 had $P \leq 0.001$. Exact topological identity was not required. This result indicates that the strongest interleaved cases retain reproducible cross-group information after pruning studies to shared species.

#### 1.6 Catalog of flagged taxa and construction of partial constraints

When evaluating the taxonomy or using it to guide tree inference, we omitted the flagged taxa's exact monophyly requirements, a procedure termed *targeted exclusion*.

For any analysis-specific species set, the raw constraint consists of the nested monophyletic descendant sets induced by represented NCBI groups. A targeted-exclusion constraint omits the monophyly requirement for a taxon only when its exact identifier occurs in the catalog of flagged taxa. All species remain in the analysis. No new clade is inserted. All unselected ancestor and descendant groups remain, and all retained groups must be nested or disjoint. The resulting object is a multifurcating partial constraint rather than a fully specified topology.

Sparse sampling can make two different NCBI nodes induce the same represented split. If an excluded taxon's descendant set is also implied by an unselected node, that split remains. This is intentional: the evidence removes one exact taxonomic claim, not every coincident split created by a particular sample.

#### 1.7 Direct validation against published trees

##### 1.7.1 Contradiction-aware normalized Robinson-Foulds distance

For partial constraint $C$ and reference tree $R$ on identical tips, let $S_C$ and $S_R$ be their unrooted nontrivial split sets. Splits $A \mid A^c$ and $B \mid B^c$ cross when all four intersections are nonempty. We defined

$$\text{c-nRF}(C,R) = \frac{|\{s \in S_C \setminus S_R : s \text{ crosses a split in } S_R\}| + |\{s \in S_R \setminus S_C : s \text{ crosses a split in } S_C\}|}{|S_C| + |S_R|}. \quad (11)$$

An unresolved constraint is therefore not penalized for failing to contain a reference split; only incompatible assertions contribute. Complete NCBI and targeted-exclusion constraints were independently restricted to the uniquely mapped species in each reference.

Among 13 recent phylogenomic reference trees, 12 were affected by exclusion of at least one flagged taxon's monophyly requirement. For each of the six reference publications also represented in the discovery collection, we used the catalog fitted to the nine cross-validation folds excluding the fold containing that publication. Every tree from the reference publication, together with the other publications in its fold, was therefore excluded from catalog selection. References absent from the discovery collection were evaluated using the full catalog. Targeted exclusion lowered c-nRF for 11 references and raised it slightly for one. The publication-balanced mean complete-NCBI-minus-targeted-exclusion change was 0.04358 (95% bootstrap interval 0.02496-0.06486; exact one-sided sign-flip $P = 0.000488$). Pooled c-nRF decreased from 0.16400 to 0.12348.

##### 1.7.2 Random-exclusion controls

Excluding the monophyly requirements of taxonomic groups can lower contradiction. We generated 10,000 sham catalogs for each of three controls: matching each targeted group on taxonomic rank, NCBI depth, $\log_2$ represented bipartition size, and informative-publication count; matching on represented size alone; or sampling uniformly. Candidate groups were represented in the reference comparison, met the corresponding publication-count filter, and were not selected by the full or held-out catalog used for that reference. Each selected taxonomic identifier was suppressed by the same constraint-building procedure used for the catalog of flagged taxa. We verified that suppressing the targeted identifiers reconstructed the exact split sets produced by the primary constraint builder.

The targeted mean c-nRF improvement was 0.04358. Fully matched, size-matched, and uniform sham catalogs had mean improvements of 0.00100, 0.00088, and 0.00281, respectively. Their 95% randomization intervals were 0.00037-0.00156, −0.00128-0.00353, and −0.00131-0.01280. None of the 10,000 replicates in any control equaled or exceeded the targeted result (one-count empirical $P = 0.0001$ for each control). Main Figure 3B plots the distribution across the 12 affected references of $(c\text{-nRF}_{\text{raw}} - c\text{-nRF}_{\text{modified}})/c\text{-nRF}_{\text{raw}}$, expressed as a percentage. Each point is one reference; random-control values are averaged over its 10,000 sham catalogs before plotting. The displayed violins therefore summarize variation among references rather than the distribution of simulation replicates.

##### 1.7.3 Publication-blocked tenfold cross-validation

We tested whether conflicts learned from one set of publications predicted contradictions in publications that did not contribute to catalog selection. Publication identifiers were ordered by a deterministic hash with seed 20260816 and assigned round-robin to ten folds. Every tree file from a publication remained in the same fold. Of the 5,865 study trees, 4,555 contained at least four uniquely mapped species and could be compared topologically.

For each fold, all publication-level observations from its publications were removed. The empirical-Bayes background, its cross-fitted sampling adjustment, group recurrence tests, adaptive publication-count filter, and BH correction were then fitted using only the other nine folds. The generic sampling-detection model described above was specified before fold assignment and reused because it had been trained on downsampling experiments rather than the focal recurrence outcomes. Thus, the cross-validation blocks the outcomes that select flagged taxa, although it is not a wholly external validation of every nuisance component. Six training sets selected a five-publication minimum and four selected a four-publication minimum; their catalogs contained 351-466 flagged taxa.

Each fold-specific catalog was evaluated only against trees from its withheld publications. Tree-level c-nRF changes were averaged within publication. Confidence intervals were obtained by resampling the affected publications, and the one-sided Monte Carlo sign-flip test used each publication's mean change as one unit with 99,999 sign reversals. The fold-level sign test treated the ten fold means as ten units.

The training-only catalogs removed at least one effective represented split from the NCBI-derived tree for 858 trees from 825 withheld publications. Targeted exclusion reduced c-nRF in 654 publications, tied in 103, and increased it in 68. The mean publication-level reduction was 0.06885 (95% publication-bootstrap interval 0.06237-0.07560; one-sided sign-flip $P = 10^{-5}$). Pooled c-nRF fell from 0.21876 to 0.19612. All ten folds had positive mean effects (one-sided sign test $P = 0.00098$).

We also tested whether this held-out improvement could be produced merely by removing the same number of comparably situated resolved relationships from each withheld tree's taxonomic representation. Within every affected tree, the control removed the same number of effective raw NCBI splits as the training-fold catalog. The true targeted splits were excluded, and replacements were chosen without replacement by nearest-neighbor matching on the taxonomic rank, NCBI depth, $\log_2$ represented split size, and informative-publication count of the NCBI group inducing each split. Controls were selected separately within each tree. Thus, unlike the coherent sham catalogs used for the phylogenomic references, this was a direct test of the mechanical scoring consequence of excluding the same number of comparably situated effective represented splits, not a test of one coherent alternative catalog. Thirty-two of the 858 affected trees contained too few nontarget splits to construct a distinct control of the same size and were excluded. Across the remaining 826 trees from 794 publications, the targeted mean publication-level improvement was 0.06212, whereas 1,000 matched-random replicates had mean improvement 0.00161 (95% randomization interval 0.00129-0.00194; one-count empirical $P = 0.000999$). The matched controls were not concentrated immediately below the discovery threshold: across the audited draws their median training-fold $q$ value was 1.0, with 0.56%, 3.18%, and 4.52% at $q \leq 0.05$, 0.10, and 0.20, respectively.

#### 1.8 Literature review of published monophyly challenges

We evaluated the 516 flagged taxa against 516 control taxa with at least five informative discovery publications and no observed contradiction in those trees. The control sample approximated the distribution of informative-publication counts in the flagged cohort. Both cohorts had a median of eight informative publications and a range of 5-37. A third sample of 516 taxa was selected without conditioning on contradiction status and is included descriptively in Supplementary Dataset 1. It overlaps 59 flagged taxa and 46 controls and was not treated as an independent third group in the inferential comparison.

OpenAlex provided candidate publications for agent-assisted review of available abstracts and selected primary full texts (Priem et al., 2022). Complete author abstracts reproduced in bibliographic records were accepted after publication-identity checks; generated summaries, mismatched text, and truncated abstracts were excluded as sources of complete abstracts. A journal article or review qualified when its examined text explicitly challenged the focal taxon's monophyly, presented an incompatible phylogenetic result, or reassessed its circumscription on that basis. Conditional findings and historical circumscriptions were retained with their qualifications. Keyword co-occurrence, weak support or unresolved placement alone, passive repetition, and synonymization without a focal monophyly challenge did not qualify. Challenges confined to constituent species, subgroups, neighboring taxa, or homonymous names were excluded. Preprints, theses, datasets, and conference abstracts were ineligible.

Each distinct publication counted once per taxon. Unassessable candidate papers were excluded individually, while each taxon remained in its original cohort and retained any qualifying evidence from other papers. Qualifying reports were matched to the 5,865-tree discovery index using PMID, DOI, normalized title, and checks against incomplete author-year identities. Reports from any publication supplying a discovery tree were removed, as were reports whose publication identity remained unresolved. The retained reports therefore had no identified overlap with discovery publications. Incomplete bibliographic identities limit the certainty of exclusion for publications without an exact match.

The primary endpoint was the fraction of taxa with at least one retained challenge report. We used 20,000 bootstrap resamples of whole taxonomic regions, defined by order, with class or phylum fallback. Both cohorts in a region were resampled together. A two-sided Fisher exact test supplied an unadjusted comparison. For a sensitivity analysis, a binomial logistic model used flagged status, $\log(1+\text{informative-publication count})$, and taxonomic rank as predictors. Ranks with fewer than 20 taxa were combined into an *other* category. Standard errors were clustered by the same taxonomic regions and a two-sided Wald test assessed flagged status. The bootstrap and regression random seed was 20261101. Region-based uncertainty accounts for within-region clustering but not every dependence introduced by papers covering several regions.

After exclusions, 231 of 516 flagged taxa (44.8%) had at least one report, compared with 64 of 516 controls (12.4%), a ratio of 3.61 (Main Fig. 4). These comprised 512 and 107 distinct taxon-publication reports, respectively. The bootstrap interval for the difference was 27.11-39.51 percentage points, and the nominal Fisher $P$ value was $1.36 \times 10^{-31}$. The adjusted odds ratio was 6.15 (95% region-clustered interval 4.36-8.68, $P = 4.19 \times 10^{-25}$). In the overlapping random sample, 125 of 516 taxa (24.2%) had a retained report. These quantities describe the yield of verified challenges in the reviewed material. A taxon without a retained report may have unexamined evidence, and some retained challenges concern historical rather than current circumscriptions.

Supplementary Dataset 1 contains original cohort membership, informative-publication counts, per-taxon report counts, bibliographic references, recorded evidence and qualifications, publication-overlap decisions, and unassessable or unresolved records. All taxa remained in their original cohorts after publication-level exclusions.

#### 1.9 Matched maximum-likelihood inference experiments

##### 1.9.1 Molecular matrices and treatment matching

Molecular matrices were assembled from GenBank records with the high-throughput workflow of Allard et al. (2026). In brief, all usable records assigned to a target clade were retrieved; putatively homologous loci were clustered with phylotaR (Bennett et al., 2018); organelle clusters were revised using genes annotated in canonical complete organelle genomes; and one sequence per species and locus was selected after sequence and locus quality control. Protein-coding markers were aligned by codon under their applicable standard, mitochondrial, or plastid genetic code, other markers were aligned as nucleotides, and excessively gappy columns and low-occupancy loci were removed.

The eligible experiments necessarily emphasize well-studied clades because they require repeated literature evidence, a recent phylogenomic reference tree, and a public sequence matrix with overlapping species. We compared them with 3,392 accepted current matrices identified from an explicit source manifest. The broader matrices had medians of 7,803 aligned sites and 10 loci. The 16 complete source matrices used in the experiments averaged 72,667 aligned sites and 105 loci; their mean site count was approximately the 94th percentile of the broader distribution. Targets of 7,735 and 1,500 sites corresponded approximately to the 50th and 10th percentiles, while targets of nine and three loci corresponded approximately to the 50th and 23rd percentiles (Supplementary Fig. S4). This enrichment motivated explicit reductions of both locus count and alignment length.

For each inference case, we used the catalog fitted after excluding the cross-validation fold containing its reference publication, including re-estimation of the background, independent filtering, and BH correction on the remaining nine folds. Each source matrix, both constraint trees, and its reference tree were then restricted to the species shared by the matrix and reference. This produced 23 reference-clade cases from 14 publications and 16 source matrices. The raw NCBI, publication-held-out targeted-exclusion, and unconstrained IQ-TREE treatments used the same taxa, alignment partitions, substitution model, random seed, and search effort. IQ-TREE 2 (Nguyen et al., 2015; Minh et al., 2020) used GTR+F+R4, `-ninit 10`, `-nbest 2`, `-safe`, and automatic threading capped at eight cores. Thus, within a replicate, the constraint treatment was the only experimental difference.

##### 1.9.2 Reduction of molecular information

We generated two independently sampled replicates at each of two reduced-data levels. Here an observed nucleotide means an unambiguous A, C, G, or T. Each reduction first chose a greedy core of loci sufficient to retain at least 20 observed nucleotides for every species, added randomly ordered loci to the requested limit, and sampled alignment columns in proportion to locus length. If the sampled columns left a species below 20 observed nucleotides, additional columns were restored until the minimum was met. The requested combinations were nine loci and 7,735 sites or three loci and 1,500 sites. Coverage protection could retain more loci than requested when no smaller set covered every species. At the 1,500-site target, the 23 cases retained one to five loci (median three), 666-1,500 aligned sites (median 1,500), and a median of 1,121 observed nucleotides per species; the least-covered species in a case had a median of 372 observed nucleotides.

##### 1.9.3 Scoring and publication-level inference

Inferred and reference trees were mutually pruned before c-nRF was calculated. For each case and replicate, improvement was the comparator c-nRF minus the targeted-exclusion c-nRF, so positive values favor targeted exclusion. Replicates were averaged within a case. Cases from the same publication were combined using their numbers of nontrivial splits in the pruned reference trees as weights. The resulting 14 publication-level effects contributed equally to the overall mean. The 95% intervals came from 100,000 bootstrap samples of whole publications, and the two-sided exact $P$ value enumerated all $2^{14}$ reversals of the signs of the publication-level effects. The bootstrap seed was 20261107, with publications ordered by identifier.

With full matrices, targeted exclusion reduced c-nRF relative to complete NCBI constraints by 0.04285 (95% publication-bootstrap interval 0.01324-0.07724; exact $P = 0.0146$). Corresponding improvements were 0.03705 at the 7,735-site target (interval 0.01204-0.06687; $P = 0.0117$) and 0.03361 at the 1,500-site target (interval 0.00799-0.06599; $P = 0.0254$). Main Figure 5A shows these mean effects, and Supplementary Figure S1 shows the underlying publication effects.

##### 1.9.4 Comparison with unconstrained inference on attainable, non-imposed reference relationships

A complete phylogenomic reference tree can contain relationships that a retained NCBI constraint makes impossible. Those relationships cannot be recovered under that constraint. Conversely, a reference split already present in the targeted-exclusion constraint should not be used to demonstrate that the remaining constraints help infer relationships from sequence data. We therefore constructed a partial reference topology in two steps. First, we removed every nontrivial reference split that crossed any split in the complete NCBI constraint. Second, we removed every remaining reference split explicitly present in the targeted-exclusion constraint. The resulting evaluation set contained only relationships that all three inference treatments could recover and that targeted exclusion did not hard-code.

For each inferred tree, we calculated c-nRF against this partial reference split set. Candidate-only and reference-only splits contributed to the numerator only when they crossed at least one split in the other set, and the denominator was the number of candidate splits plus the number of retained reference splits. All inference treatments for a case contained the same species and the same number of candidate splits. Replicates were averaged within case. Cases were weighted by their numbers of retained reference splits when combined within publication. The resulting publication-level effects contributed equally to the overall mean, with confidence intervals and exact tests calculated as above.

Targeted exclusion reduced c-nRF relative to no constraint by 0.02076 with full data (95% publication-bootstrap interval 0.00611-0.03944; exact $P = 0.0117$), 0.05655 at the 7,735-site target (interval 0.02131-0.09301; $P = 0.0107$), and 0.10465 at the 1,500-site target (interval 0.06437-0.14803; $P = 0.00037$). Publication counts favoring targeted exclusion, tying, or favoring no constraint were 8/5/1, 10/1/3, and 13/0/1, respectively. These results are shown in Main Figure 5B.

##### 1.9.5 Reference-tree-size-weighted sensitivity analysis

As a separate sensitivity analysis, we weighted publication-level effects by the total number of scored reference splits in each publication. Bootstrap sampling and exact sign reversals still operated on complete publications. Relative to complete NCBI constraints, the weighted improvements were 0.04345 with full data (95% interval 0.00916-0.08864; $P = 0.0371$), 0.03222 at 7,735 sites (0.00530-0.07113; $P = 0.0547$), and 0.03150 at 1,500 sites (0.00385-0.07475; $P = 0.0693$). For the non-imposed, NCBI-compatible reference relationships, weighted improvements over no constraint were 0.02947 (0.01052-0.04593; $P = 0.0195$), 0.04114 (0.02216-0.06769; $P = 0.00488$), and 0.06624 (0.03686-0.10898; $P = 0.00098$), respectively. Both weighting schemes therefore gave positive mean effects at all three data levels, although the weighted tests against complete NCBI constraints did not reach 0.05 at the reduced-data levels.

##### 1.9.6 Dependence on disagreement between the reference tree and NCBI

For each case, taxonomic disagreement was the fraction of nontrivial reference splits that crossed at least one split in the complete NCBI constraint. For publications represented by multiple cases, conflicting and total reference-split counts were summed before the fraction was calculated. We then tested the association between this publication-level disagreement and the c-nRF advantage of targeted exclusion over no constraint. Spearman rank correlations were evaluated by 99,999 two-sided permutations of the publication effects.

When all reference splits were scored, targeted exclusion became less favorable as disagreement with complete NCBI increased. Spearman $\rho$ was $-0.722$ with full data ($P = 0.00486$), $-0.723$ at the 7,735-site target ($P = 0.00458$), and $-0.682$ at the 1,500-site target ($P = 0.00945$). After incompatible reference splits were excluded, disagreement no longer predicted the targeted-exclusion advantage: absolute $\rho$ ranged from 0.13 to 0.34 and all $P \geq 0.231$. Only one of 14 publications had no incompatible reference split. For descriptive stratification, that publication was therefore combined with the seven nonzero publications at or below the median nonzero disagreement (0.1693); the remaining six formed the higher-disagreement group (Supplementary Fig. S2).

##### 1.9.7 Limitations of the inference experiments

Restricting matrices to species shared with a phylogenomic reference does not reproduce every feature of inference in a larger species set, because additional low-data taxa can interact with focal placements. Each reduced matrix was sampled twice, and IQ-TREE searches were not repeated under multiple random seeds. The plotted $P$ values were not adjusted across site targets. The partial-reference endpoint deliberately asks how well the methods recover relationships that remain attainable under all constraint treatments; it does not measure recovery of reference relationships forbidden by retained constraints. Finally, a published phylogenomic tree is a high-quality reference rather than known truth, and c-nRF gives the same topological weight to every resolved reference split regardless of branch duration. The comparisons therefore estimate performance in the sampled clades under the specified species restrictions and scoring rules.

#### 1.10 Upper confidence limits on recurring contradiction

Failure to detect recurring contradiction does not establish that a group is monophyletic: the available publications may simply provide too little evidence. We therefore also calculated an upper confidence limit for the recurrence parameter $\rho_g$, using the same publication outcomes and fitted background and detection probabilities as the primary analysis. A low upper limit indicates that the observed record is inconsistent with a strong recurring tendency to contradict the group, under the fitted model. It is not a probability that the group is monophyletic.

For each group, we held $f_{ig}$ and $D_{ig}$ fixed and used $p_{ig}(r) = f_{ig} + (1 - f_{ig})D_{ig}r$ for each proposed recurrence value $r \in [0, 1]$. The log likelihood was

$$\ell_g(r) = \sum_i \{Y_{ig} \log p_{ig}(r) + (1 - Y_{ig}) \log[1 - p_{ig}(r)]\}.$$

We compared each proposed value with the best-fitting value using $T_g(r) = 2[\ell_g(\widehat{\rho}_g) - \ell_g(r)]$. Confidence sets were obtained by Neyman test inversion with likelihood-ratio ordering, following the general construction of Feldman and Cousins (1998), adapted here to publications with different Bernoulli probabilities rather than their Poisson examples. For each proposed value, we calculated how often a publication record generated at that value would produce a likelihood-ratio statistic at least as large as the observed statistic. Proposed values with tail probability greater than 0.05 were retained. The reported limit is the largest retained value: the upper endpoint of an individual conditional 95% confidence set, not an asymptotic one-sided 95% interval.

For groups with at most 12 informative publications, all $2^n$ possible binary publication records were enumerated. Each record was weighted by its probability at the proposed recurrence value, and its own best-fitting recurrence was used in its likelihood-ratio statistic. For larger records, we simulated 4,000 records at each proposed value, increasing to

20,000 when the preliminary tail probability was within 0.015 of 0.05. Monte Carlo probabilities included the observed record through the usual plus-one correction. Ties were included in the upper tail. Random streams were fixed by taxonomic identifier and proposed recurrence value so that resuming the calculation did not change results.

If recurrence equal to one was retained, the upper limit was one. Otherwise, we evaluated the full interval on a grid spaced by 0.025, including the fitted recurrence, and refined the interval above the highest accepted grid value at spacing 0.0025. Finite upper limits were padded upward by one fine-grid increment and capped at one. Thus, reported endpoints are numerical approximations, with Monte Carlo uncertainty for larger publication records. Exact enumeration checks confirmed at least 95% conditional coverage for the small-record designs examined, and independent simulation checks assessed larger designs. These checks concern the confidence-set construction, not uncertainty in the learned background and detection models.

Limits were calculated for all 18,109 groups with informative publication records, including groups below the five-publication filter used for discovery. The remaining 8,182 of the 26,291 represented groups had no informative publication record and were assigned missing limits rather than interpreted as supported. Of the calculated limits, 8,134 were below one and 9,975 equaled one (Supplementary Fig. S6). The resource includes the fitted recurrence, its upper limit, publication counts, background and detection summaries, and calculation method. These are individual, conditional limits: they assume the fitted probabilities and independent publication outcomes, do not propagate uncertainty in model training or unknown publication dependence, and do not provide simultaneous confidence guarantees across taxa. No threshold on these limits was used to select a new catalog or alter the 516 flagged taxa.

##### 1.10.1 Optional externally supplied study weights

The software additionally accepts positive, prespecified publication weights based on criteria such as data quantity or recency. No such metadata-based weights were used in this study. The reported analyses use background and detection probabilities estimated from the study trees.

For the optional extension, the supplied weights are normalized to mean one within each focal group and multiply the publication contributions to $\ell_g(r)$. The weighted likelihood-ratio statistic is calibrated by simulating publication outcomes under the same null model and refitting with the same fixed weights. Weights do not increase publication counts or change the replication filter. Background and detection fitting remain unchanged. Weighted runs omit the asymptotic computational screen and bootstrap all observed groups. Weights must be specified independently of focal outcomes. This software option has unit tests, including exact agreement with the default when weights are uniform, but has not undergone the end-to-end validation reported for the unweighted analysis. The upper confidence limits above use the original, unweighted likelihood.

#### 1.11 Exploratory analysis of higher taxonomic ranks

The primary analysis focused on families and lower ranks to assess fine-scale taxonomic groupings. To examine the applicability of the workflow beyond that scope, we separately analyzed ranked groups above family through phylum using the same 5,865 study trees from 5,072 source publications and the 22 July 2026 NCBI taxonomy snapshot. We retained the original frozen tip mappings: only uniquely mapped species represented once in a tree contributed to group classification. The extension included superfamily, parvorder, infraorder, suborder, order, superorder, subcohort, cohort, infraclass, subclass, class, superclass, subphylum, and phylum. Named unranked groups above family were not included in this extension.

Background and detection models were fitted anew using the higher-rank observations, rather than applying probabilities learned for families and lower ranks. The detection model used 2,000 contradicted tree-group templates, with 40 downsampled records and one complete record per template (82,000 training records). Cross-fitting held out broad taxonomic regions, and prediction used the corresponding held-out model for observations from represented training regions or the final fitted model for previously unseen regions. The workflow used five folds, a 201-value recurrence grid, 99,999 parametric-bootstrap replicates per computationally screened group, and the same broad asymptotic screening threshold of 0.30. The informative-publication filter was selected from cutoffs of 2-20, and BH adjustment was applied jointly across all higher ranks passing the selected filter. No externally supplied study weights were used.

Of 3,123 represented groups, 876 met the selected five-publication filter and 55 had adjusted $q \leq 0.05$ (Supplementary Fig. S7). These included 21 orders, 11 suborders, nine superfamilies, four subclasses, three classes, two infraorders, two phyla, and one each at cohort, parvorder, and superorder. The two flagged phyla were Methanobacteriota and Bacillota. The complete evidence table and a separate list of the 55 flagged taxa are supplied with the higher-rank results. These groups are reported as a separate exploratory resource.

The template-weighted held-out Brier score for detection was 0.2133, compared with 0.3665 when every detection probability was set to one. The observed-prevalence constant gave a descriptive Brier score of 0.2322, but was not a separately fold-trained comparator. The background model's cross-fitted Brier score was 0.1138. These diagnostics assess prediction within the extension. End-to-end null calibration and independent literature or inference validation were not repeated for higher ranks, so these results are exploratory and the nominal BH threshold should not be interpreted as a separately demonstrated 5% false-discovery guarantee for this extension. Differences among rank-specific flagged fractions also reflect sampling and nested membership and are not estimates of the prevalence of incorrect taxonomy at those ranks.

#### 1.12 AI use acknowledgement

Agentic AI tools, including OpenAI Codex, assisted with developing and implementing statistical methods, writing and debugging analysis code, searching and evaluating the literature, preparing figures, and editing the manuscript. The authors take responsibility for the methods, results, citations, and final text.

#### 1.13 Supplementary datasets

| Item | Contents |
|---|---|
| Supplementary Dataset 1 | Taxonomic-literature review: cohort membership, per-taxon counts, publication references, evidence assessments, qualifications, publication-overlap decisions, and unassessable or unresolved records. |
| All-group evidence | All 26,291 represented NCBI taxa, publication counts, displayed and contradicting counts, fitted recurrence, bootstrap *P* value, independent-filter status, BH *q* value, and action. |
| Publication observations | The 73,181 informative assessments of individual NCBI taxa within publications, including classification, sample sizes, fitted background probability, and detection probability. |
| Flagged taxa | The 516 exact NCBI identifiers flagged for recurrent contradiction of monophyly, with names, ranks, evidence summaries, *P* values, and *q* values. |
| Topology-recurrence evidence | The 132 shared-species topology-recurrence cases, contributing publications, information statistics, permutation distributions, and Monte Carlo *P* values. |

| Item | Contents |
|---|---|
| Reference-tree provenance | Provenance, taxon mapping, molecular-data scale, and citations for phylogenomic reference trees. |
| Validation comparisons | c-nRF comparisons with phylogenomic references, coherent matched sham catalogs, and publication-blocked tenfold cross-validation results for TimeTree study trees. |
| Inference manifest | Publication-held-out IQ-TREE run manifest, molecular matrix, retained reduction treatment, constraint treatment, software settings, and c-nRF. |
| Upper-limit resource | All 26,291 represented groups, including individual conditional 95% recurrence upper limits for 18,109 groups and explicit missing-evidence status for 8,182 groups. |
| Higher-rank resource | Evidence for 3,123 ranked groups above family through phylum, the separate list of 55 flagged groups, rank-level summaries, run settings, and model diagnostics. Exploratory extension, not an addition to the primary catalog. |

### 2 Supplementary figures

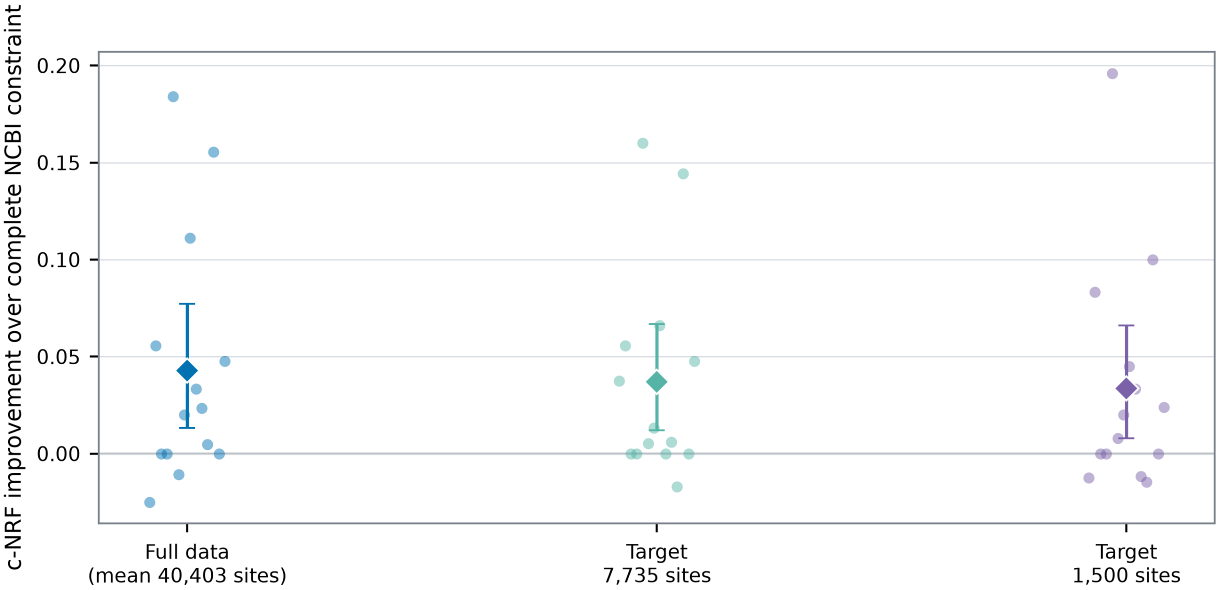


Figure S1: **Publication-level effects underlying Main Figure 5A.** Small points show the c-nRF improvement from targeted exclusion relative to the complete NCBI constraint for individual publications. Diamonds show means giving each publication equal weight, and error bars show 95% intervals from resampling whole publications. Full matrices contained a mean of 40,403 aligned sites after restriction to reference species; reduced matrices targeted 7,735 or 1,500 sites across all species. Positive values favor targeted exclusion.

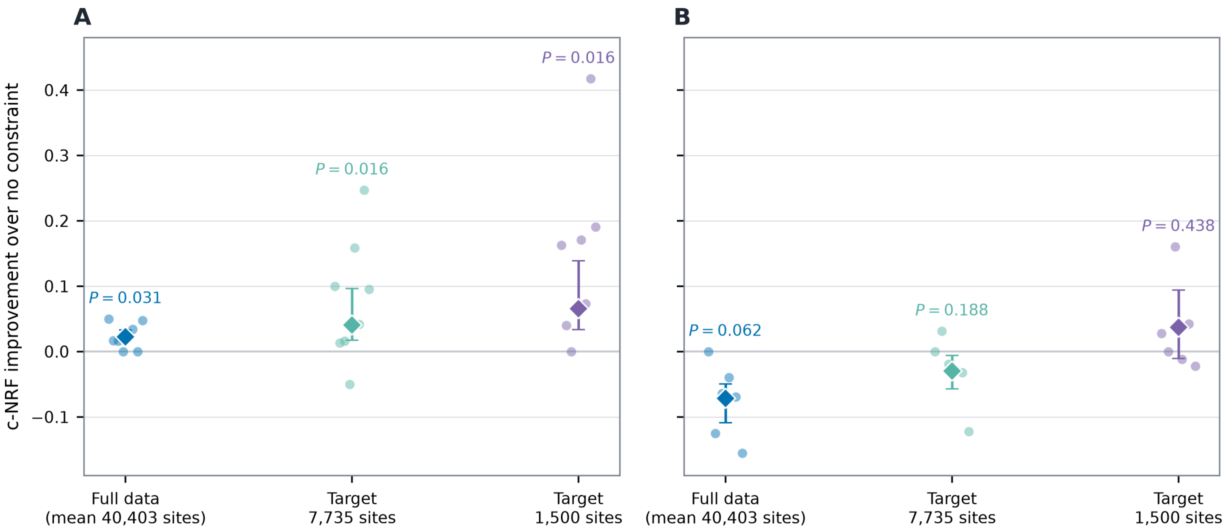


Figure S2: **Size-weighted sensitivity analysis stratified by disagreement between the phylogenomic reference and complete NCBI.** The 23 clade-level comparisons were combined within their 14 source publications before plotting and statistical analysis. Values use c-nRF against the complete reference tree, before incompatible reference relationships are removed. Positive values favor targeted exclusion over no constraint. **(A)** The single publication with no incompatible reference split and the seven publications at or below the median nonzero disagreement. **(B)** The six publications above that median. Small points show individual publication effects. Diamonds show means weighted by the number of nontrivial reference splits, error bars show 95% intervals from resampling whole publications, and labels give two-sided exact publication-level sign-reversal tests. The continuous correlation analysis is primary because only one publication had zero disagreement.

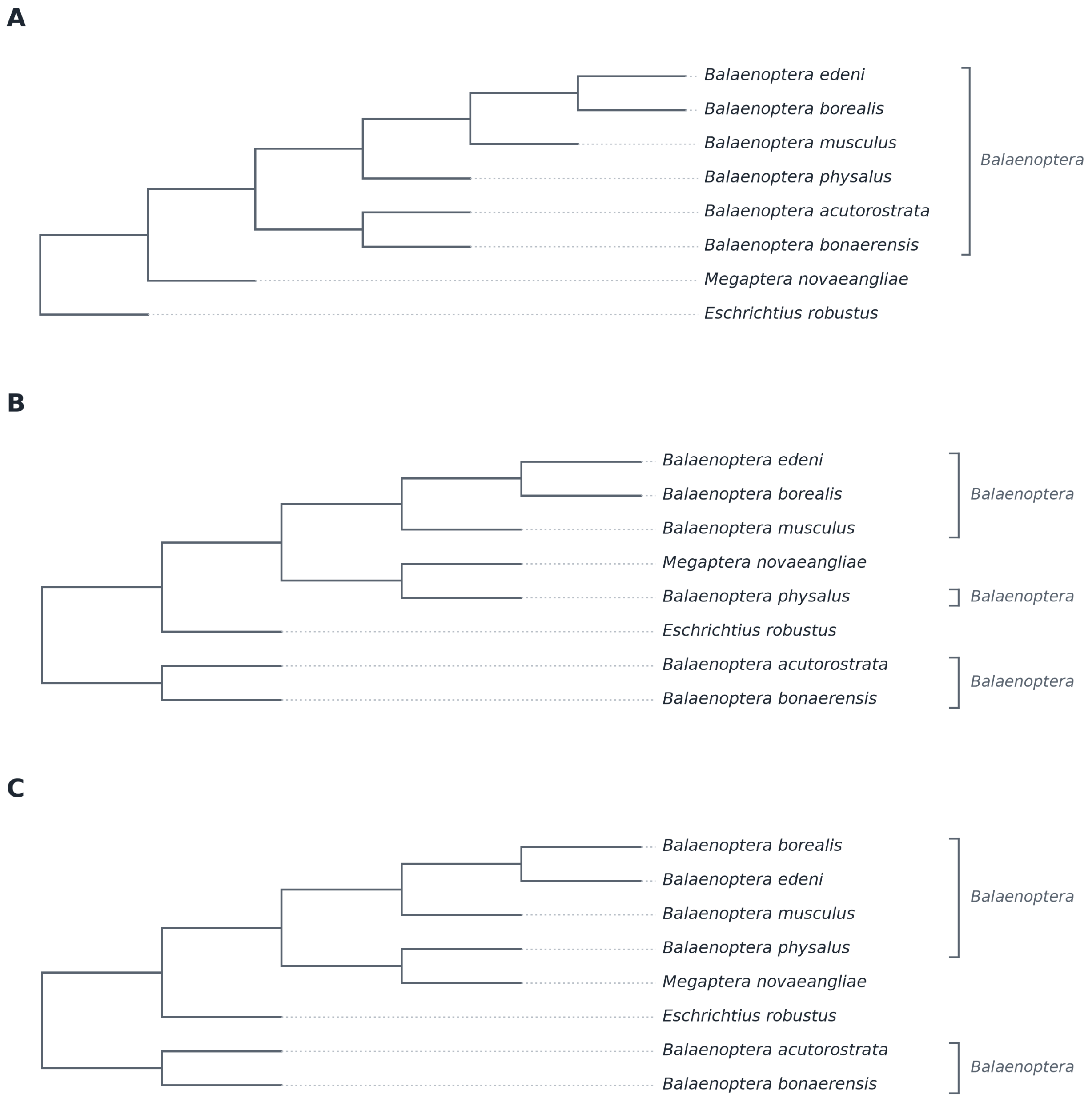


Figure S3: **Baleen-whale positive control.** Trees inferred from the same cetacean matrix and pruned to eight whales under **(A)** complete NCBI-derived constraints (three of five focal splits matched) and **(B)** targeted exclusion of the contradicted *Balaenoptera* and Balaenopteridae constraints (five of five matched), compared with **(C)** the 3,191-gene ASTRAL phylogenomic reference tree of McGowen et al. (2020). Trees are cladograms; horizontal lengths have no scale.

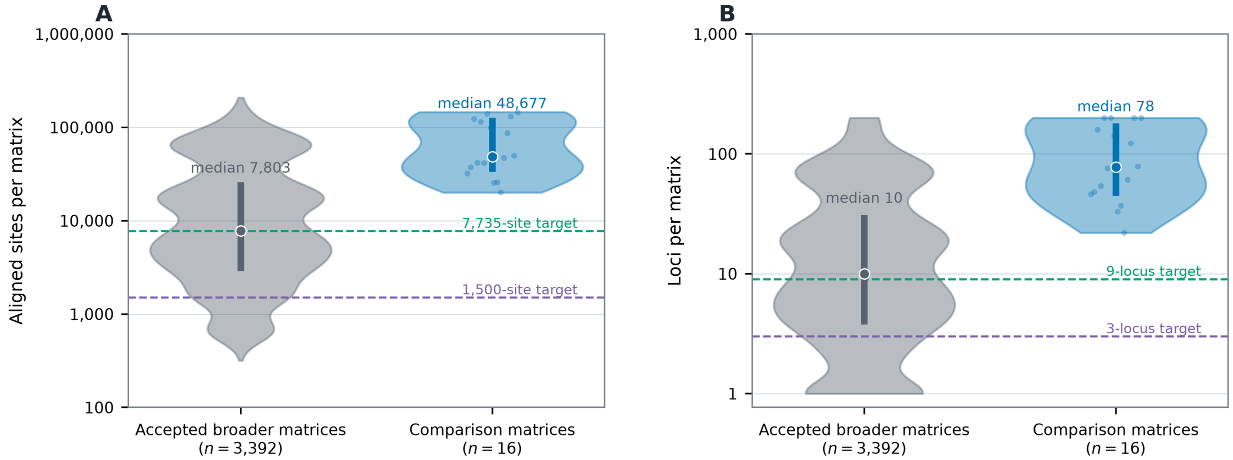


Figure S4: **Data scale of the inference experiments.** **(A)** Aligned sites and **(B)** loci in 3,392 accepted current matrices identified from the complete source manifest and the 16 complete source matrices used for inference experiments. Violin widths show distributions, thick segments show interquartile ranges, circles show medians, and small points show individual comparison matrices; axes are logarithmic. The broader matrices had medians of 7,803 sites and 10 loci, whereas the comparison matrices averaged 72,667 sites and 105 loci. In the broader distribution, 7,735 and 1,500 sites lie at approximately the 50th and 10th percentiles, while nine and three loci lie at approximately the 50th and 23rd percentiles. Dashed lines mark these targets.

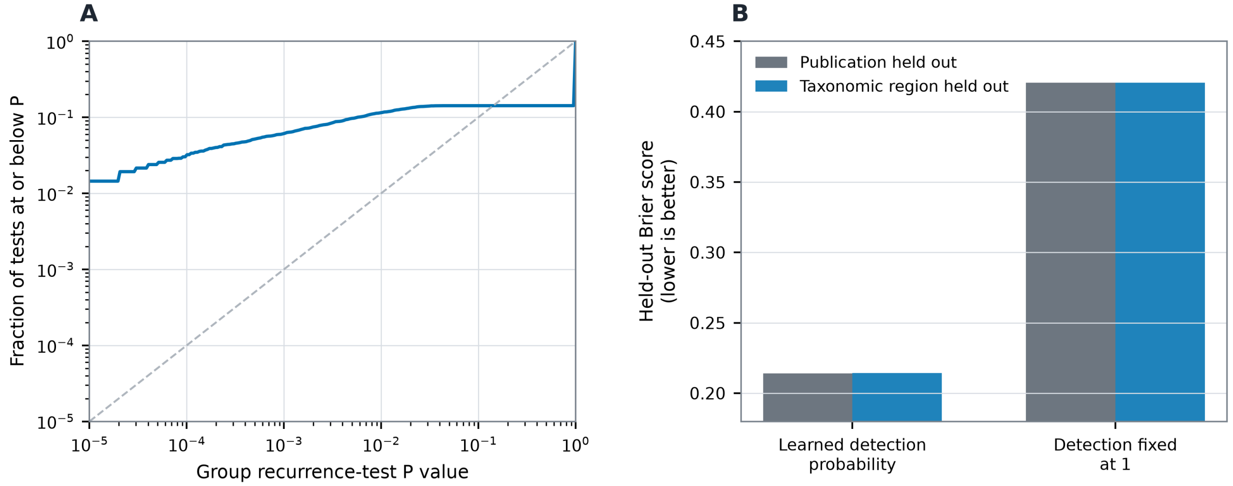


Figure S5: **Statistical diagnostics for recurrence testing and sampling detection.** **(A)** Cumulative distribution of bootstrap $P$ values for the 5,395 groups meeting the five-publication filter. Groups excluded by the broad asymptotic computational screen were assigned $P = 1$ before BH correction. **(B)** Held-out Brier scores for the learned sampling-detection model and the ablation that set every detection probability to one. Scores use 80,000 reduced-tree observations derived from 2,000 contradicted complete-tree templates, with equal total weight per template. Complete publications or complete broad taxonomic regions were held out during model fitting; lower values indicate better prediction.

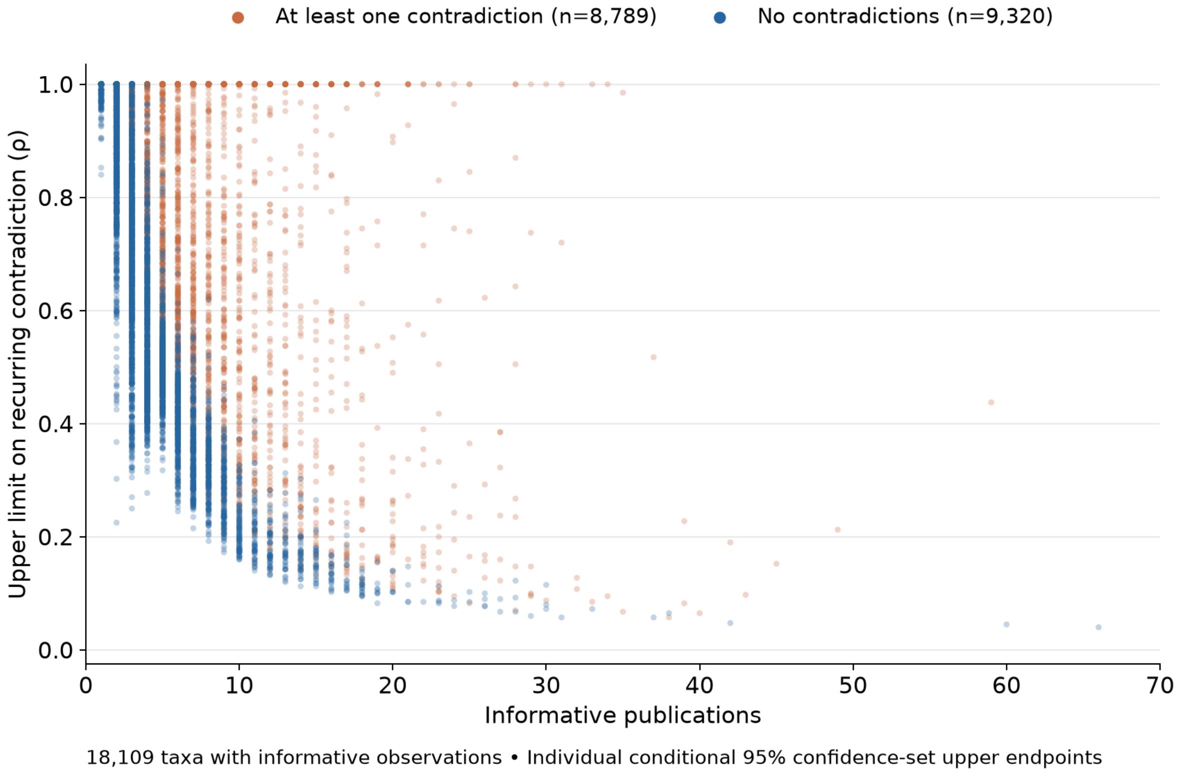


Figure S6: **Upper confidence limits on recurring contradiction.** Each point represents an NCBI group with an informative publication record. The horizontal axis gives the number of informative publications, and the vertical axis gives the upper endpoint of its individual conditional 95% confidence set for recurrence, $\rho$. Orange points have at least one observed contradiction and blue points have none. Points are transparent and may overlap. Low limits require sufficiently informative evidence against high recurrence, not merely absence from the catalog of flagged taxa. A limit of one means that maximal recurrence could not be excluded. All 18,109 groups with informative records are shown. The 8,182 groups without informative publications have no calculated limit and are omitted. Limits condition on the learned background and detection probabilities and are not simultaneous confidence statements or probabilities of monophyly (see Supplementary Methods, Section 1.10).

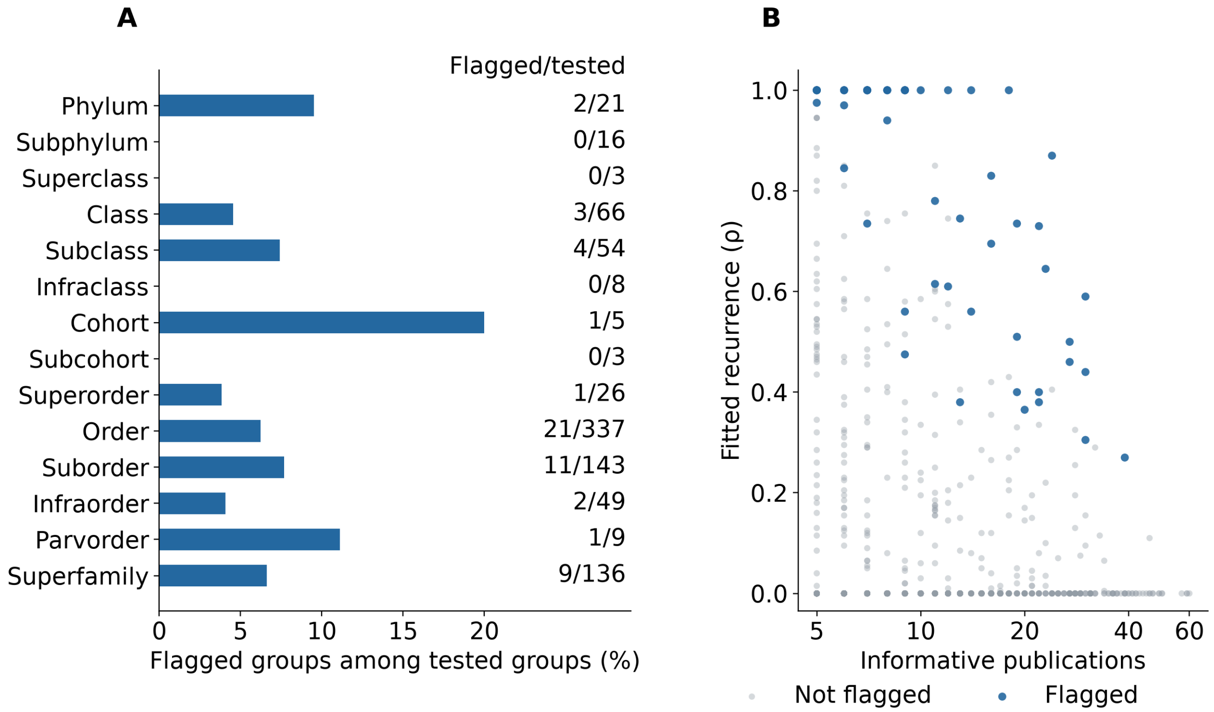


Figure S7: **Exploratory recurrence analysis above family through phylum. (A)** Percentage of tested groups flagged at nominal BH $q \leq 0.05$, ordered from higher to lower ranks. Labels give flagged/tested group counts. The analysis represented 3,123 groups, of which 876 met the selected five-publication filter and 55 were flagged. Fractions are descriptive and can be based on small denominators. **(B)** Fitted recurrence versus informative-publication count for the 876 tested groups. Blue points are flagged groups and gray points are other tested groups; transparency shows overlapping observations. A large fitted recurrence alone does not imply a significant result. Background and detection models were retrained for this separate higher-rank analysis, but the primary analysis's end-to-end calibration and external validations were not repeated (see Supplementary Methods, Section 1.11).

### References


John Allard, Jack Craig, and Sudhir Kumar. Completing a timetree of eukaryotes with high-throughput phylogenetics. Manuscript in preparation, 2026.

Yoav Benjamini and Yosef Hochberg. Controlling the false discovery rate: a practical and powerful approach to multiple testing. *Journal of the Royal Statistical Society, Series B*, 57(1):289-300, 1995. doi:10.1111/j.2517-6161.1995.tb02031.x.

Yoav Benjamini and Daniel Yekutieli. The control of the false discovery rate in multiple testing under dependency. *The Annals of Statistics*, 29(4):1165-1188, 2001. doi:10.1214/aos/1013699998.

Dominic J. Bennett, Hannes Hettling, Daniele Silvestro, Alexander Zizka, Christine D. Bacon, Søren Faurby, Rutger A. Vos, and Alexandre Antonelli. phylotaR: An automated pipeline for retrieving orthologous DNA sequences from GenBank in R. *Life*, 8(2):20, 2018. doi:10.3390/life8020020.

Richard Bourgon, Robert Gentleman, and Wolfgang Huber. Independent filtering increases detection power for high-throughput experiments. *Proceedings of the National Academy of Sciences of the United States of America*, 107(21): 9546-9551, 2010. doi:10.1073/pnas.0914005107.

Gary J. Feldman and Robert D. Cousins. Unified approach to the classical statistical analysis of small signals. *Physical Review D*, 57:3873-3889, 1998. doi:10.1103/PhysRevD.57.3873.

Jerome H. Friedman. Greedy function approximation: a gradient boosting machine. *The Annals of Statistics*, 29(5): 1189-1232, 2001. doi:10.1214/aos/1013203451.

S. Blair Hedges, Joel Dudley, and Sudhir Kumar. TimeTree: a public knowledge-base of divergence times among organisms. *Bioinformatics*, 22(23):2971-2972, 2006. doi:10.1093/bioinformatics/btl505.

Nikolaos Ignatiadis, Bernd Klaus, Judith B. Zaugg, and Wolfgang Huber. Data-driven hypothesis weighting increases detection power in genome-scale multiple testing. *Nature Methods*, 13:577-580, 2016. doi:10.1038/nmeth.3885.

Sangjin Kim and Paul Schliekelman. Prioritizing hypothesis tests for high throughput data. *Bioinformatics*, 32(6): 850-858, 2016. doi:10.1093/bioinformatics/btv608.

Sudhir Kumar, Glen Stecher, Michael Suleski, and S. Blair Hedges. TimeTree: a resource for timelines, timetrees, and divergence times. *Molecular Biology and Evolution*, 34(7):1812-1819, 2017. doi:10.1093/molbev/msx116.

Sudhir Kumar, Michael Suleski, Jack M. Craig, et al. TimeTree 5: an expanded resource for species divergence times. *Molecular Biology and Evolution*, 39(8):msac174, 2022. doi:10.1093/molbev/msac174.

Michael I. Love, Wolfgang Huber, and Simon Anders. Moderated estimation of fold change and dispersion for RNA-seq data with DESeq2. *Genome Biology*, 15:550, 2014. doi:10.1186/s13059-014-0550-8.

Michael R. McGowen, Georgia Tsagkogeorga, Sandra Álvarez-Carretero, et al. Phylogenomic resolution of the cetacean tree of life using target sequence capture. *Systematic Biology*, 69(3):479-501, 2020. doi:10.1093/sysbio/syz068.

Bui Quang Minh, Heiko A. Schmidt, Olga Chernomor, et al. IQ-TREE 2: new models and efficient methods for phylogenetic inference in the genomic era. *Molecular Biology and Evolution*, 37(5):1530-1534, 2020. doi:10.1093/molbev/msaa015.

Carl N. Morris. Parametric empirical Bayes inference: theory and applications. *Journal of the American Statistical Association*, 78(381):47-55, 1983. doi:10.1080/01621459.1983.10477920.

Lam-Tung Nguyen, Heiko A. Schmidt, Arndt von Haeseler, and Bui Quang Minh. IQ-TREE: a fast and effective stochastic algorithm for estimating maximum-likelihood phylogenies. *Molecular Biology and Evolution*, 32(1): 268-274, 2015. doi:10.1093/molbev/msu300.

Fabian Pedregosa, Gaël Varoquaux, Alexandre Gramfort, Vincent Michel, Bertrand Thirion, Olivier Grisel, Mathieu Blondel, Peter Prettenhofer, Ron Weiss, Vincent Dubourg, Jake Vanderplas, Alexandre Passos, David Cournapeau, Matthieu Brucher, Matthieu Perrot, and Édouard Duchesnay. Scikit-learn: machine learning in Python. *Journal of Machine Learning Research*, 12:2825-2830, 2011.

Belinda Phipson and Gordon K. Smyth. Permutation *P*-values should never be zero: calculating exact *P*-values when permutations are randomly drawn. *Statistical Applications in Genetics and Molecular Biology*, 9(1):Article 39, 2010. doi:10.2202/1544-6115.1585.

Jason Priem, Heather Piwowar, and Richard Orr. OpenAlex: A fully-open index of scholarly works, authors, venues, institutions, and concepts, 2022. arXiv:2205.01833.

Steven G. Self and Kung-Yee Liang. Asymptotic properties of maximum likelihood estimators and likelihood ratio tests under nonstandard conditions. *Journal of the American Statistical Association*, 82(398):605-610, 1987. doi:10.1080/01621459.1987.10478472.

Martin R. Smith. Information theoretic generalized robinson-foulds metrics for comparing phylogenetic trees. *Bioinformatics*, 36(20):5007-5013, 2020. doi:10.1093/bioinformatics/btaa614.

Mervyn Stone. Cross-validatory choice and assessment of statistical predictions. *Journal of the Royal Statistical Society, Series B*, 36(2):111-133, 1974. doi:10.1111/j.2517-6161.1974.tb00994.x.
